# Environmental stochasticity can account for patterns of within-host respiratory virus evolution

**DOI:** 10.64898/2026.05.15.725410

**Authors:** Wenfei Fiona Xiao, Mireille Farjo, Anice Lowen, Katia Koelle

## Abstract

The ecological and evolutionary dynamics of populations, including viral populations, are known to be jointly shaped by deterministic and stochastic processes. While the impact of stochastic processes has been rigorously explored for viral dynamics at the level of the host population, most dynamic models for acutely-infecting respiratory viral pathogens at the within-host scale remain deterministic in their formulation. While this may be reasonable for identifying key processes shaping their within-host viral population dynamics, recent studies indicate that stochastic processes need to be invoked for understanding patterns of within-host viral evolution. Specifically, several studies have shown that viral allele frequencies can change dramatically over the time course of days in acute infections. Here, we use stochastic dynamic models to explore the role of environmental noise in shaping observed patterns of virus evolution in acute respiratory virus infections. We summarize ways in which environmental stochasticity can be biologically realized in these acute viral infections and describe within-host models that can be implemented to jointly yield viral population dynamics and evolutionary dynamics. We further develop a statistical approach to estimate the extent of environmental noise from observed within-host allele frequency changes. We test this approach on simulated data and apply it to existing influenza A virus and SARS-CoV-2 within-host data. With these applications, we show that environmental stochasticity can parsimoniously reproduce key features of empirically observed allele frequency changes without needing to invoke demographic stochasticity or to adopt Wright-Fisher model formulations with a constant effective population size. Finally, we show that purifying selection and positive selection can both still contribute to within-host viral evolution in the context of a noisy environment, providing theoretical support for studies that have found purifying and positive selection in acutely-infecting respiratory virus populations.

## Introduction

Viral pathogens reproduce and evolve across multiple biological scales. At the within-host scale, various mechanistic models have been fit to viral titer data to capture and better understand the population dynamics of acutely-infecting respiratory viral pathogens (Baccam et al., 2006; Saenz et al., 2010; Smith and Perelson, 2011; Pawelek et al., 2012; Ke et al., 2021). Other studies have shown that the genetic diversity of these viral populations are typically very low (Murcia et al., 2010; Dinis et al., 2016; McCrone et al., 2018), likely as a result of stringent transmission bottlenecks (McCrone et al., 2018; Martin and Koelle, 2021). Moreover, the frequencies of detected intrahost Single Nucleotide Variants (iSNVs) have been shown to change rapidly over the time course of hours to days (McCrone et al., 2018; VanInsberghe et al., 2024). This pattern is similarly observed for nonsynonymous and synonymous iSNVs and has lead to the conclusion that stochastic processes are the dominant drivers of within-host viral evolution (McCrone et al., 2018). The presence of purifying selection has also been consistently detected in these viral populations (Dinis et al., 2016; McCrone et al., 2018; VanInsberghe et al., 2024). In contrast, positive selection has not been readily detected in natural infections of acutely infecting respiratory viruses, although it has been detected in experimental challenge studies (Sobel Leonard et al., 2016; Ferreri et al., 2026), including those involving non-human animal models Wilker et al. (2013); Plante et al. (2021); Zhou et al. (2021).

More broadly, stochastic processes are known to structure communities and populations and to generate variability in nature (Coulson et al., 2004; Shoemaker et al., 2020). Two distinct types of stochastic processes have been characterized in the literature: demographic stochasticity and environmental stochasticity. Demographic stochasticity describes the realized variability that arises from the probabilistic nature of demographic processes such as births and deaths (Lande, 1993). While this type of stochasticity is important when population sizes are small, demographic stochasticity becomes a negligible source of process noise when population sizes are large. Environmental stochasticity describes realized variability that arises from extrinsic variation in environmental conditions that affect population growth rates and survival (Fujiwara and Takada, 2017). This type of stochasticity remains an important source of noise even when population sizes are large. While mathematical models can include both of these sources of stochasticity, they commonly include only one. Models that include demographic stochasticity often are implemented using the exact or approximate Gillespie algorithm (Gillespie, 1977, 2001). However, Wright-Fisher models with constant population sizes also implement demographic stochasticity, with genetic drift becoming more pervasive at small effective population sizes. Models that instead include environmental stochasticity are often implemented using the Euler-Maruyama simulation method (Kloeden and Platen, 1992a), where extrinsic variation in environmental conditions is captured by adding a stochastic term to the model that introduces noise on top of a deterministic skeleton.

Recent studies have applied Wright-Fisher models to estimate within-host effective viral population sizes (*N*_*E*_ values) for human influenza viruses using observed iSNV frequency changes from longitudinally sampled infected individuals (McCrone et al., 2020; Lumby et al., 2020; Shi et al., 2024; Ferreri et al., 2026). While a study that analyzed a chronic influenza infection estimated a large *N*_*E*_ value of 2.5 × 10^7^, studies examining within-host viral evolution in acute infections have arrived at values that are extremely small, on the order of 20-70 viruses (McCrone et al., 2020; Shi et al., 2024; Ferreri et al., 2026). These low *N*_*E*_ estimates reflect the large changes in iSNV frequencies that have been empirically observed between samples taken 1-3 days apart. However, within-host viral population sizes are generally very large, such that models with small effective viral population sizes become mechanistically less plausible, even when considering substantial heterogeneity in viral output across infected cells (Russell et al., 2018). Moreover, census viral population sizes *N* vary by orders of magnitude over the course of an acute infection. Because 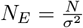, a constant *N*_*E*_ would require parallel, orders-of-magnitude changes in the variance of the offspring distribution *σ*^2^ to compensate for this change in *N*. Such parallel changes in the variance of the offspring distribution seem unlikely to us. Together, these considerations indicate to us that a Wright-Fisher model may not appropriately capture the most relevant source of noise shaping viral allele frequency changes within acute viral infections. More generally, they indicate to us that models assuming that changes in allele frequencies result from demographic stochasticity may be limited in terms of biological plausibility, although their ability to reproduce certain features of observed changes in allele frequencies has been demonstrated.

Here, we therefore ask whether environmental stochasticity may instead be able to account for observed patterns of within-host viral evolution in acute respiratory infections. Environmental noise is reasonable to expect in within-host viral dynamics, with extrinsic variation in environmental conditions likely to impact the rate at which viral particles infect target cells and the rate at which virus is cleared, among other processes governing viral growth and survival (Table 1). To quantitatively consider environmental stochasticity as a driver of within-host viral evolution, we first present mechanistic and phenomenological within-host models that incorporate environmental noise and then show how these models can be extended to account for viral genetic variation and within-host viral evolution. We then derive a statistical inference approach for estimating the extent of environmental noise from observed within-host iSNV frequency changes and test it on simulated data. Applying this approach to a previously analyzed human influenza A virus (IAV) dataset (McCrone et al., 2018) and to a previously analyzed SARS-CoV-2 dataset (Tonkin-Hill et al., 2021), we find that environmental stochasticity can account for observed allele frequency changes and that the model yields predictions that are consistent with observed evolutionary patterns. Finally, using model simulations, we examine the efficiency of within-host purifying selection and positive selection in the context of viral evolution shaped by environmental stochasticity.

**Table 1.**
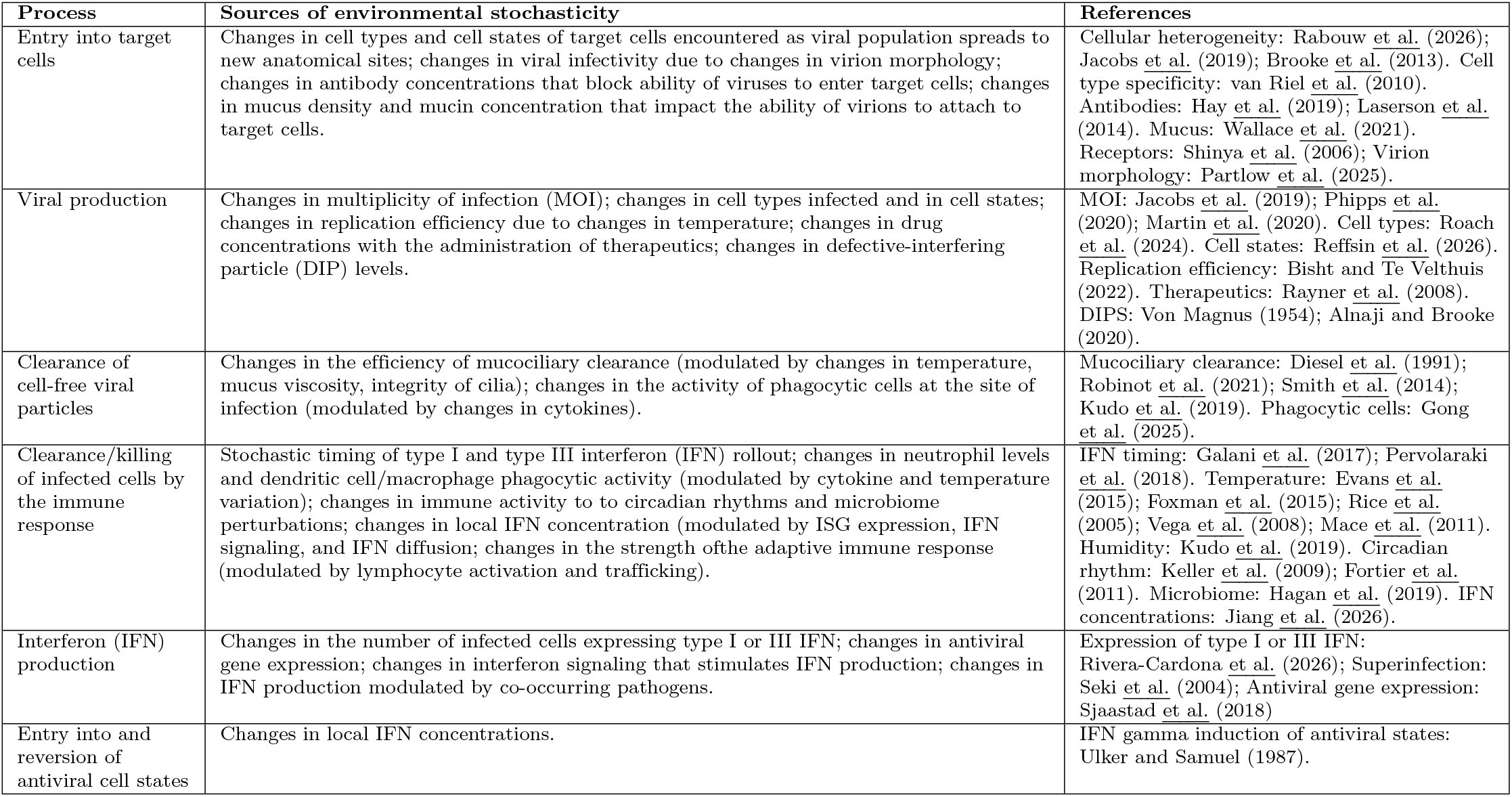
Processes impacting within-host viral dynamics that can be affected by variation in environmental conditio. lists a process that impacts within-host viral dynamics and briefly describes what changes would result in variation in environment Supporting references are provided in the third column.

| Process | Sources of environmental stochasticity | References |
| --- | --- | --- |
| Entry into target cells | Changes in cell types and cell states of target cells encountered as viral population spreads to new anatomical sites; changes in viral infectivity due to changes in virion morphology; changes in antibody concentrations that block ability of viruses to enter target cells; changes in mucus density and mucin concentration that impact the ability of virions to attach to target cells. | Cellular heterogeneity: Rabouw et al. (2026); Jacobs et al. (2019); Brooke et al. (2013). Cell type specificity: van Riel et al. (2010). Antibodies: Hay et al. (2019); Laserson et al. (2014). Mucus: Wallace et al. (2021). Receptors: Shinya et al. (2006); Virion morphology: Partlow et al. (2025). |
| Viral production | Changes in multiplicity of infection (MOI); changes in cell types infected and in cell states; changes in replication efficiency due to changes in temperature; changes in drug concentrations with the administration of therapeutics; changes in defective-interfering particle (DIP) levels. | MOI: Jacobs et al. (2019); Phipps et al. (2020); Martin et al. (2020). Cell types: Roach et al. (2024). Cell states: Reffsin et al. (2026). Replication efficiency: Bisht and Te Velhuis (2022). Therapeutics: Rayner et al. (2008). DIPs: Von Magnus (1954); Alnaji and Brooke (2020). |
| Clearance of cell-free viral particles | Changes in the efficiency of mucociliary clearance (modulated by changes in temperature, mucus viscosity, integrity of cilia); changes in the activity of phagocytic cells at the site of infection (modulated by changes in cytokines). | Mucociliary clearance: Diesel et al. (1991); Robinot et al. (2021); Smith et al. (2014); Kudo et al. (2019). Phagocytic cells: Gong et al. (2025). |
| Clearance/killing of infected cells by the immune response | Stochastic timing of type I and type III interferon (IFN) rollout; changes in neutrophil levels and dendritic cell/macrophage phagocytic activity (modulated by cytokine and temperature variation); changes in immune activity to circadian rhythms and microbiome perturbations; changes in local IFN concentration (modulated by ISG expression, IFN signaling, and IFN diffusion; changes in the strength of the adaptive immune response (modulated by lymphocyte activation and trafficking). | IFN timing: Galani et al. (2017); Pervolaraki et al. (2018). Temperature: Evans et al. (2015); Foxman et al. (2015); Rice et al. (2005); Vega et al. (2008); Mace et al. (2011). Humidity: Kudo et al. (2019). Circadian rhythm: Keller et al. (2009); Fortier et al. (2011). Microbiome: Hagan et al. (2019). IFN concentrations: Jiang et al. (2026). |
| Interferon (IFN) production | Changes in the number of infected cells expressing type I or III IFN; changes in antiviral gene expression; changes in interferon signaling that stimulates IFN production; changes in IFN production modulated by co-occurring pathogens. | Expression of type I or III IFN: Rivera-Cardona et al. (2026); Superinfection: Seki et al. (2004); Antiviral gene expression: Sjaastad et al. (2018) |
| Entry into and reversion of antiviral cell states | Changes in local IFN concentrations. | IFN gamma induction of antiviral states: Ulker and Samuel (1987). |

## Methods

### Stochastic within-host viral population dynamics

We start with a mechanistic model that describes the within-host viral dynamics of an acute respiratory infection. Originally developed for influenza A virus infections, the model incorporates the roles of the innate and the adaptive immune response in regulating within-host viral dynamics and has been fit to a combination of viral titer data and interferon data (Pawelek et al., 2012). The deterministic model equations are: 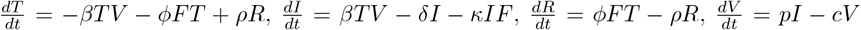, and 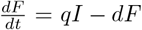, where *T* are uninfected target cells, *I* are infected target cells, *R* are uninfected target cells that are refractory to infection, *V* is cell-free virus, and *F* are interferon levels. Uninfected target cells become infected at rate *βTV* and become refractory to infection at rate *ϕFT* through exposure to interferon. Refractory cells lose their protection from infection at rate *ρR*. Infected cells die at rate *δI*, with *δ* = *δ*_*I*_ prior to the onset of the adaptive immune response at time *t*_*µ*_ and *δ* = *δ*_*A*_ after time *t*_*µ*_. They are also removed at rate *κIF*, which implements the role of innate immune response cells such as natural killer cells in infected cell death. Infected cells produce free virus at rate *pI* and free virus is cleared at rate *cV*. Finally, interferon is produced by infected cells at rate *qI* and interferon decay occurs at rate *dF*. Environmental stochasticity can be incorporated into this model by adding noise into one or more of the modeled processes and numerically simulating the model using the Euler-Maruyama approach (Kloeden and Platen, 1992b). Incorporating noise into viral clearance yields the following set of equations:

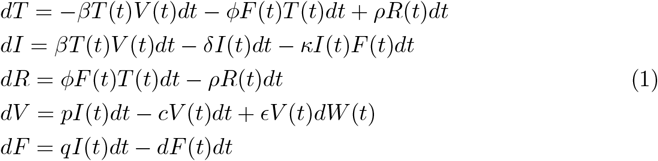

where *ϵ* (≥ 0) quantifies the degree of environmental noise and *dW* (*t*) is a normally distributed random variables with mean 0 and variance *dt*.

In addition to this mechanistic within-host model, we can use a simple piecewise exponential phenomenological model to capture the general shape of within-host viral dynamics. Models of this form have been successfully fit to viral titer data from various viral infections, most recently SARS-CoV-2 (Kissler et al., 2021a,b). This phenomenological model contains four parameters: the proliferation rate *b*, peak viral titer, time of peak viral titer *τ*_*p*_, and the clearance rate *c*. When *t < τ*_*p*_, the viral population is exponentially growing, with 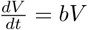. When *t > τ*_*p*_, the viral population is exponentially declining, with 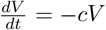. Incorporating environmental noise, these equations can be written as:

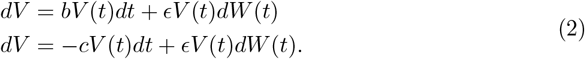

### Stochastic viral dynamics and evolution

We can extend the above model to consider a viral population containing an intrahost Single Nucleotide Variant (iSNV). In this case, we have two viral ‘strains’, which we can call *V*_1_ and *V*_2_. We (for now) assume that these strains differ from one another genetically but not phenotypically. For the piecewise exponential model, during viral proliferation, the dynamics of the viral strains are given by:

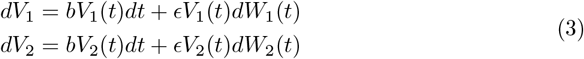

Similarly, during viral clearance, the dynamics of the viral strains are given by:

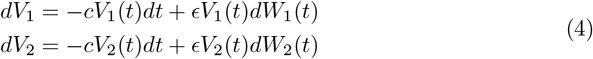

During both viral proliferation and viral clearance, we allow for the possibility of correlation between the environmental noise impacting the two strains, with parameter *ρ* (0 ≤ *ρ* ≤ 1) quantifying the degree of correlation. Such correlation could arise as a result of sharing the same spatial environment within the respiratory tract. Under this conceptualization, lower correlations could thus reflect spatial compartmentalization of the two strains, even within the sampled nasal environment. With the possibility of noise correlation, at each time step, the noise values *dW*_1_(*t*) and *dW*_2_(*t*) are therefore drawn from multivariate normal distributions with means of 0 and a covariance matrix given by 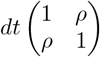. We are interested in the evolutionary dynamics of the two strains in the viral population. At any time *t*, the frequency of strain 2 can be calculated as:

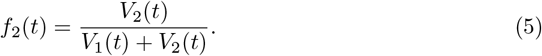

The frequency of strain 1 at time *t* is then simply *f*_1_(*t*) = 1 − *f*_2_(*t*).

### Derivation of a model that can be used for statistical inference

We seek an analytical expression for *f*_2_ that does not explicitly involve viral titers *V*_1_ and *V*_2_ such that we can estimate parameters of a model using only allele frequency data, rather than both allele frequency data and viral titer data. Towards this goal, we define:

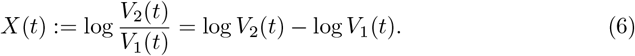

With this definition, the frequency of strain 2 is given by:

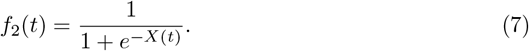

In the Appendix, we show that allele frequency changes of strain 2 under the dynamic models given by equations 3 and 4 are dynamically equivalent to those of a Brownian motion model of *X*(*t*) given by:

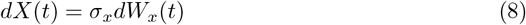

where *dW*_*x*_(*t*) is white noise, drawn from a normal distribution with mean 0 and variance *dt*. Called the diffusion parameter in Brownian motion models, the parameter *σ*_*x*_ quantifies the intensity of the random fluctuations and is a function of *ρ* and *ϵ*:

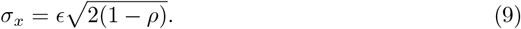

We thus have a model that governs the dynamics of within-host viral evolution that is independent of viral titers. From equation 9, it can be seen that the intensity of the random fluctuations depends only on two parameters: the magnitude of environmental noise *ϵ* and the degree of correlation *ρ* between the environmental noise experienced by the two strains. At a given level of environmental noise *ϵ*, we expect allele frequencies to change most dramatically when the correlation of the environmental noise is lower (*ρ* closer to 0). With full correlation of environmental noise (*ρ* = 1), the frequencies of the two strains each stay constant, regardless of the amount of environmental noise. At a given degree of noise correlation *ρ*, we expect allele frequencies to change more dramatically when the magnitude of environmental noise *ϵ* is larger. When *ϵ* = 0, there is no noise and the frequencies of the two strains each stay constant.

### Simulation of mock datasets

We forward-simulated allele frequency dynamics under three different *σ*_*x*_ parameterizations (*σ*_*x*_ = 0.6, *σ*_*x*_ = 1.0, and *σ*_*x*_ = 1.4) and used these generated mock datasets to determine the extent to which we could recover the *σ*_*x*_ values used in the forward simulations. To simulate these iSNV dynamics, we simulated the Brownian motion model for *X*(*t*) and converted the simulated *X*(*t*) values into allele frequencies *f*_2_(*t*) using equation 7. Fifty iSNV trajectories were generated for each *σ*_*x*_ value considered, each representing frequency dynamics of a single iSNV from a single infected individual. For all of the simulated trajectories, the initial iSNV frequency was drawn from an exponential distribution with a mean of 0.15. iSNVs with initial frequencies below a variant calling threshold of 2% were discarded and replaced with another randomly drawn initial iSNV frequency. We generated 3 mock datasets for each *σ*_*x*_ value considered. The first mock dataset assumed that samples were taken from infected individuals on sequential days (one day apart). The second and third mock datasets assumed that samples were instead taken 2 days and 4 days apart, respectively. For each dataset, the first observation time point was the same and we defined this time point as time *t* = 0.

### Estimation of diffusion parameter *σ*_*x*_ from simulated mock datasets

To determine the extent to which we could recover *σ*_*x*_ from each of these mock datasets, we converted the allele frequencies at times *t* = 0 and *t* = *T* (where *T* = 1, 2, and 4 days for the three different datasets) to their corresponding *X*(*t*) values using equation 7. To estimate *σ*_*x*_, we make use of analytical expressions derived for Brownian motion models, namely that *X*(*t* = *T*) is distributed according to a normal distribution with mean *X*(*t* = 0) and variance 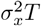. For each paired-time iSNV observation, we can thus calculate the probability of observing *X*(*t* = *T*) given *X*(*t* = 0), a value for *σ*_*x*_, and the value of *T*. Because variant-calling thresholds lead to censored data points, iSNVs with frequencies at time *t* = *T* that fall below the variant-calling threshold or above one minus the variant-calling threshold need to be handled slightly differently. The probability of observing an iSNV that falls below the variant-calling threshold (here, of 2%) at time *t* = *T* is given by a normal cumulative distribution function evaluated at 2%. The probability of observing an iSNV that falls above a frequency of 98% at time *t* = *T* is given by one minus the normal cumulative distribution function evaluated at 98%. These normal cumulative distribution functions have a mean of *X*(*t* = 0) and a variance of 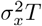. For a given *σ*_*x*_, once we have a probability evaluated for each paired-time iSNV observation, we sum the logs of these probabilities over all observations, yielding the log-likelihood of the Brownian motion model parameterized with the given *σ*_*x*_ value.

### Analysis of empirical datasets

We analyzed two previously published deep-sequencing viral datasets of longitudinally-sampled within-host acute respiratory infections. The first dataset we analyzed stems from a study by McCrone et al. (2018) which focused on viral evolution in influenza A virus (IAV) populations. The second dataset we analyzed stems from a study by Tonkin-Hill et al. (2021) which focused on viral evolution in SARS-CoV-2 populations. We point the reader to these previous publications for detailed information on the sequencing protocols they used, as well as information on coverage and read depth.

For the IAV dataset, we relied on a supplemental file of a recent publication (Shi et al., 2026) that estimated the within-host viral effective population size (*N*_*E*_) for the within-host IAV data from McCrone et al. (2018). In that study, *N*_*E*_ was estimated to be small (28-84 viral particles) using a beta-with-spikes approximation of the Wright-Fisher model (Tataru et al., 2015). The iSNV frequencies within this supplemental file (their Table S1) closely matched those provided in the original McCrone et al. (2018) paper. Here, we used only a subset of the paired-time iSNV observations from this supplemental file. The subset of observations used were those in which the iSNV frequency at the first time point of the pair was above the variant-calling threshold of 2%. The iSNV frequency at the second time point of the pair could be above or below 2%. Each paired-time observation we extracted contained the ID of the sampled individual (e.g., 50001), the iSNV identity (e.g., PA G271A), collection times of the first and second samples from this paired-time observation (in days post symptom onset, e.g., 0 and 6), and the frequency of the focal iSNV in the first and second samples (e.g., 0.0916 and 0). Our dataset contains a total of 60 paired-time observations from 28 unique individuals. We provide these paired-time observations in our Table S1.

For the SARS-CoV-2 dataset, we relied on iSNV frequencies estimated and made publicly available by Tonkin-Hill et al. (2021). We again limited our analyses to iSNVs where the frequency at the first sampled time point was above 2%, regardless of whether the second time point was above or below 2%. Each paired-time observation contained the ID of the sampled individual (e.g., CAMS001914), the IDs of the paired samples (e.g., CAMB-71C42 and CAMB-79DCF), the collection dates of the two samples (e.g., 3/31/2020 and 4/2/2020), the time between the two samples (in days), the iSNV identity (e.g., G24781C), and the frequencies of the focal iSNV in the first and second samples (e.g., 0.2081 and 0.0388). Only point mutation iSNVs were considered (insertions and deletions of any length were excluded). When multiple samples were collected from the same individual on the same day, the sample with the lowest Ct value (highest viral titer) was selected for analysis. We also filtered out sites documented in https://virological.org/t/issues-with-sars-cov-2-sequencing-data/473 as mutations that arise recurrently across SARS-CoV-2 lineages and are recommended for masking, which led to the removal of 11 iSNV observations. Our final SARS-CoV-2 dataset contains a total of 54 paired-time observations from 22 unique individuals. We provide these paired-time observations in our Table S2.

### Model criticism

In addition to estimating *σ*_*x*_ for the empirical IAV and SARS-CoV-2 datasets, we evaluate whether and the extent to which the Brownian motion model developed here matches observed patterns of iSNV frequency changes in these datasets. To this end, we first again note that under our Brownian motion model, the difference *X*(*t* = *T*) − *X*(*t* = 0) follows a normal distribution with mean 0 and variance 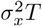. For a normal random variable with mean 0 and variance 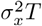, its absolute value follows a half-normal distribution with expected value given by:

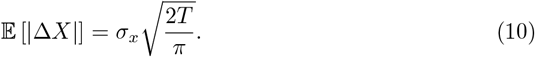

This expression provides a theoretical prediction for how the magnitude of absolute changes in *X* scales with elapsed time under our model with environmental stochasticity. We can now compare these theoretical predictions against observations from mock datasets that we simulated as well as against empirical observations. The mock datasets we generated were simulated under a Brownian motion model for *X* with *σ*_*x*_ = 1.5. Initial iSNV frequencies were, as before, drawn from an exponential distribution with a mean of 0.15 (and redrawn is the value fell below our variant calling threshold of 0.02). 50 simulated iSNV trajectories had their second observation taken at time *T* = 0.0417 days (1 hour following the first observation). Five more sets of 50 trajectories were simulated, corresponding to second observations taken at times *T* = 0.25 days (6 hours following the first observation), *T* = 0.5 days (12 hours following the first observation), *T* = 1 day, *T* = 2 days, and *T* = 3 days. Absolute changes in *X* were calculated for each of these 300 simulated trajectories.

We further calculated absolute changes in *X* for the empirical IAV and SARS-CoV-2 datasets. Because some paired-time observations had the second timepoint iSNV frequency falling below the variant-calling threshold of 2%, we calculated mean absolute changes in *X* using two different approaches. Approach (1) excluded paired-time observations that had iSNV frequencies of zero at the second time point, while retaining observations that had non-zero estimated iSNV frequencies at the second time point (regardless of whether they fell above or below the variant calling threshold of 2%). Approach (2) retained all paired-time observations but set the iSNV frequencies at the second observation time points to the variant-calling threshold of 2% if they fell below this threshold. Observed absolute changes in *X* were plotted alongside the theoretical expectation, as were the means of the observed absolute changes in *X*. Two means were plotted, corresponding to the two different approaches for handling second timepoint iSNV frequencies below-the-variant calling threshold.

Finally, we compared observed changes in *X* (that is, our Δ*X* values) against the theoretical prediction that these values should be normally distributed with mean 0 and variance 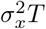. This comparison was done for both the simulated data set and for the empirical datasets. Using the one-sample Kolmogorov-Smirnov goodness-of-fit test, we determined whether each of these three Δ*X* datasets were statistically different from reference normal distributions parameterized with their estimated *σ*_*x*_ values. We used Approach (1) in handling second time point iSNVs whose frequencies were 0%.

### Assessment of purifying and positive selection

The Brownian motion model we developed above assumes that all genetic variation analyzed is neutral with respect to fitness. This Brownian motion model can be extended in a straightforward manner to consider iSNVs or variants that have either a selective advantage or a selective disadvantage. We derive this extension in the Appendix. The extended model derivation results in the frequency changes of an iSNV being given by:

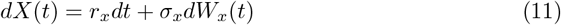

where *r*_*x*_ is defined as either *b*_2_ − *b*_1_ (if *t < τ*_*p*_) or *c*_1_ − *c*_2_ (if *t > τ*_*p*_). This model is an Arithmetic Brownian Motion model with drift term *r*_*x*_*dt* and diffusion term *σ*_*x*_*dW*_*x*_(*t*). The focal iSNV has a selective disadvantage when *r*_*x*_ *<* 0 and a selective advantage when *r*_*x*_ *>* 0. When *r*_*x*_ = 0, the iSNV is fitness neutral and equation 12 reduces to the original Brownian motion model given by equation 8. Under this model, an iSNV with initial frequency *X*(*t* = 0) is distributed at time *T* as:

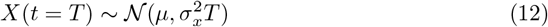

where *µ* = *X*(*t* = 0) + *r*_*x*_*T*. Using this expression, we can calculate the probability that an iSNV with frequency *X* at time *t* = 0 is detected above the variant-calling threshold at time *T*. This probability depends both on the selective effect of the iSNV mutation (*r*_*x*_) and on the parameter *σ*_*x*_ that quantifies the amount of environmental stochasticity.

## Results

### Simulations of within-host viral dynamics incorporating environmental stochasticity

Figure 1A shows simulated viral dynamics under the mechanistic within-host model given by equations 1 with parameter values for pony 1 in Pawelek et al. (2012). Three viral dynamic simulations are shown: one without environmental noise (*ϵ* = 0), one with a low level of environmental noise (*ϵ* = 0.5), and one with a higher level of environmental noise (*ϵ* = 1.0). It is evident from these simulations that environmental noise can impact viral dynamics, although the overall trajectory of the viral dynamics still reflects the processes implemented in the deterministic skeleton of the Pawelek et al. (2012) model.

**Figure 1.**
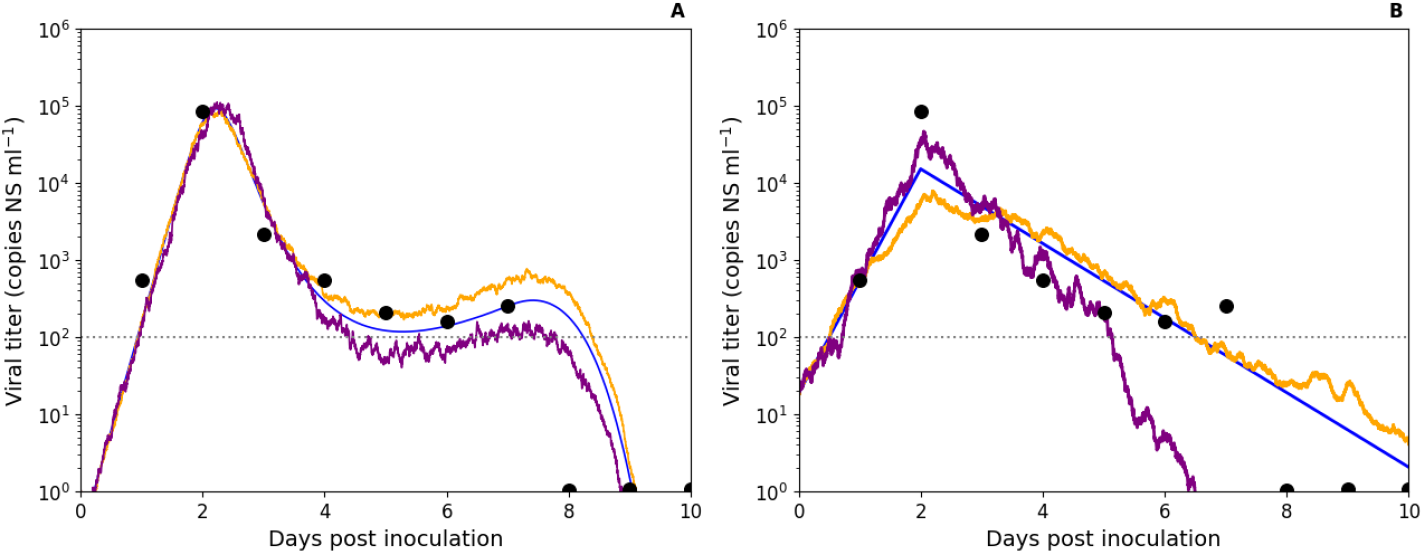
Simulated viral dynamics incorporating environmental stochasticity. (A) Stochastic simulations of the Pawelek et al. (2012) model, incorporating both innate and adaptive immune responses. The model was parameterized with pony 1 values from Table 1 and Table 2 of Pawelek et al. (2012). Filled circles show viral titer measurements for pony 1. The blue line shows a simulation without environmental noise (*ϵ* = 0.0). The orange line shows a simulation with a low amount of environmental noise (*ϵ* = 0.5). The purple line shows a simulation with a higher amount of environmental noise (*ϵ* = 1.0). (B) Stochastic simulations of the piecewise exponential model, parameterized with maximum likelihood estimates of the viral proliferation rate (*b* = 3.36 per day) and the viral clearance rate (*c* = 1.11 per day). As in panel (A), filled circles show viral titer measurements for pony 1 and three simulations with differing levels of environmental noise: *ϵ* = 0.0 (blue line), *ϵ* = 0.5 (orange line), and *ϵ* = 1.0 (purple line). In both (A) and (B), the horizontal line at 10^2^ copies NS ml^−1^ shows the assay’s limit of detection.

Figure 1B shows simulated viral dynamics under the piecewise exponential model given by equations 2 and parameterized using maximum likelihood estimates of the viral proliferation rate *b* and clearance rate *c* for pony 1’s measured viral titers. Three viral dynamic simulations are again shown: one without environmental noise (*ϵ* = 0), one with a low level of environmental noise (*ϵ* = 0.5), and one with a higher level of environmental noise (*ϵ* = 1.0). Again, it is evident from these simulations that environmental noise can impact viral dynamics, although the overall trajectory of the viral dynamics still reflect the overall processes of viral proliferation and clearance.

### Simulations of the two-strain model with environmental stochasticity

Figure 2 shows stochastic simulations of the two-strain piecewise exponential viral dynamic model, with the second strain initially present at a frequency of 20%. We simulated this model under four different parameterizations. Viral proliferation rates and viral clearance rates were the same across each parameterization, but the extent of environmental noise *ϵ* and the degree of environmental noise correlation *ρ* differed across simulations. In the first column, environmental noise was low (*ϵ* = 0.35) and the degree of environmental noise correlation was high (*ρ* = 0.95). This resulted in similar and largely deterministic viral dynamics of the two strains (Figure 2A) and only small changes in allele frequencies over the 8-day simulated time course (Figure 2B). In the second column, environmental noise remained low (*ϵ* = 0.35) but we set the degree of environmental noise correlation to be low (*ρ* = 0.05). This still resulted in largely deterministic viral dynamics of the two strains (Figure 2D) and relatively small changes in the frequency of strain 2 over the 8-day simulated time course (Figure 2E), albeit the simulated frequency changes were more pronounced than in Figure 2B. In the third column, environmental noise was higher (*ϵ* = 0.8) and we set the degree of environmental noise correlation to be high (*ρ* = 0.95). This resulted in more stochastic viral dynamics for both viral strains (Figure 2G) but only small changes in strain 2’s frequencies over the 8-day simulated time course (Figure 2H). Indeed, the frequency changes in Figure 2H are less pronounced than those observed when environmental noise was lower but the correlation in noise was higher (Figure 2E). Finally, in the fourth column, environmental noise was set to *ϵ* = 0.8 and we lowered the degree of environmental noise correlation to *ρ* = 0.05. This again resulted in visibly stochastic viral dynamics for both viral strains (Figure 2J) as well as pronounced changes in strain 2’s frequencies over the 8-day simulated time course (Figure 2K). These simulations show that both the extent of environmental noise *ϵ* and the degree of correlation between environmental noise *ρ* impact within-host allele frequency dynamics.

**Figure 2.**
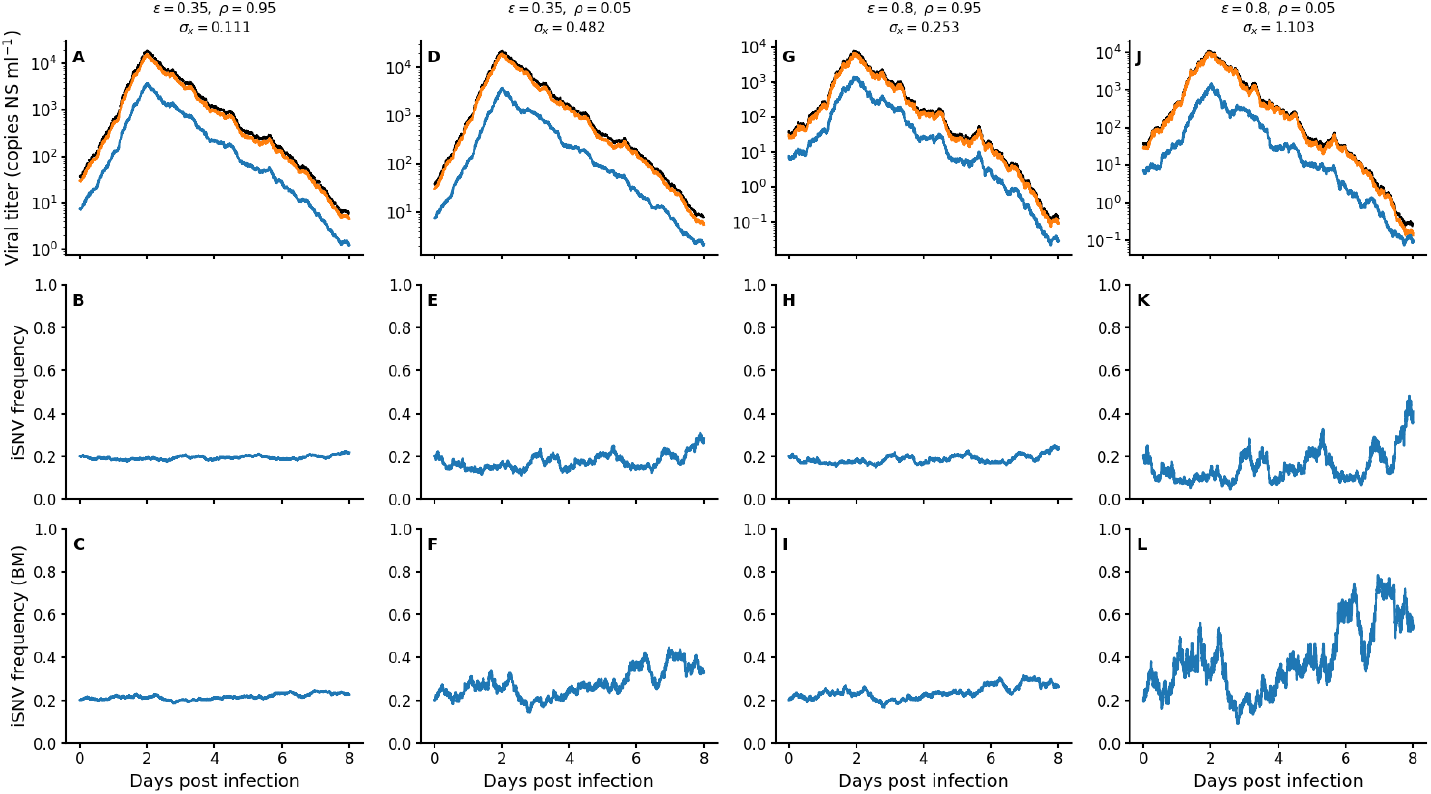
Simulated two-strain viral population dynamics and evolutionary dynamics. Columns correspond to different *ϵ* and *ρ* parameterizations. (A) Viral titers from simulations of the two-strain piecewise exponential model. Titers are shown separately for the two strains (strain 1 in orange and strain 2 in blue) as well as overall (in black, *V*_1_ + *V*_2_). Here, *ϵ* = 0.35 and *ρ* = 0.95. (B) Frequency changes of strain 2, calculated from panel (A) using equation 5. (C) Frequency changes of strain 2 from a simulation of the Brownian motion model (equation 8) with same initial strain frequency as in panel (B) and using a *σ*_*x*_ value of 0.111, calculated using *ϵ* = 0.35 and *ρ* = 0.95. (D) Viral titers from simulations of the two-strain piecewise exponential model. Here, *ϵ* = 0.35 and *ρ* = 0.05. Colors are as in panel (A). (E) Frequency changes of strain 2, calculated from panel (D). (F) Frequency changes of strain 2 from a simulation of the Brownian motion model using a *σ*_*x*_ value of 0.482, calculated using *ϵ* = 0.35 and *ρ* = 0.05. (G) Viral titers from simulations of the two-strain piecewise exponential model. Here, *ϵ* = 0.8 and *ρ* = 0.95. Colors are as in panel (A). (H) Frequency changes of strain 2, calculated from panel (G). (I) Frequency changes of strain 2 from a simulation of the Brownian motion model using a *σ*_*x*_ value of 0.253, calculated using *ϵ* = 0.8 and *ρ* = 0.95. (J) Viral titers from simulations of the two-strain piecewise exponential model. Here, *ϵ* = 0.8 and *ρ* = 0.05. Colors are as in panel (A). (K) Frequency changes of strain 2, calculated from panel (J). (L) Frequency changes of strain 2 from a simulation of the Brownian motion model using a *σ*_*x*_ value of 1.103, calculated using *ϵ* = 0.8 and *ρ* = 0.05. The two-strain piecewise exponential model in panels (A), (D), (G), and (J) was parameterized with a proliferation rate of *b* = 3.36 per day and a clearance rate of *c* = 1.11 per day.

We next performed simulations to demonstrate that our derived Brownian motion model would recapitulate simulated patterns of allele frequency dynamics when appropriately parameterized. Specifically, we calculated the *σ*_*x*_ value for each of the four scenarios described above and simulated the Brownian motion model given by equation 8 with these *σ*_*x*_ values, starting with an initial *X* value corresponding to *f*_2_ = 20%. We then converted the simulated *X* dynamics into allele frequency dynamics using equation 7. The last row of Figure 2 shows representative simulations of iSNV frequencies from this Brownian motion model. As expected, the second row simulations and the third row simulations yield dynamically similar allele frequency dynamics.

### The parameter *σ*_*x*_ can be recovered from mock datasets

We used our simulated iSNV trajectories to assess whether we could statistically recover the value of the diffusion parameter *σ*_*x*_ (quantifying the intensity of random fluctuations) from forward simulations of the Brownian motion model with a given value of this parameter. Figures 3A,B, and C each show 10 representative simulations of our mock data trajectories. The three columns correspond to three different values of *σ*_*x*_ used to generate the mock datasets: 0.6, 1.0, and 1.4. Time *t* = 0 corresponds to the time of the first collected sample. The vertical dashed lines show the times of the second collected sample (T = 1 day, 2 days, and 4 days). The second through fourth rows correspond to inference of *σ*_*x*_ for the 50 mock datasets with the second collection time point being T = 1 day, T = 2 days, and T = 4 days, respectively. Across all three considered values of *σ*_*x*_ and sampling time differences, the true value of *σ*_*x*_ was contained within the 95% confidence interval (CI) of the maximum likelihood estimate (MLE). Notably (and to us, surprisingly), the time point of the second collected sample *T* had no appreciable impact on the width of the 95% confidence interval. These results demonstrate that inference of *σ*_*x*_ is robust to variation in temporal spacing between observations within the range considered here. This result is desirable in practice, as it implies that for empirical datasets with different sampling intervals, the precision of *σ*_*x*_ estimates will not be appreciatively governed by the particular sampling scheme employed. The width of the confidence interval does scale with the magnitude of *σ*_*x*_, with higher *σ*_*x*_ values having broader confidence intervals.

**Figure 3.**
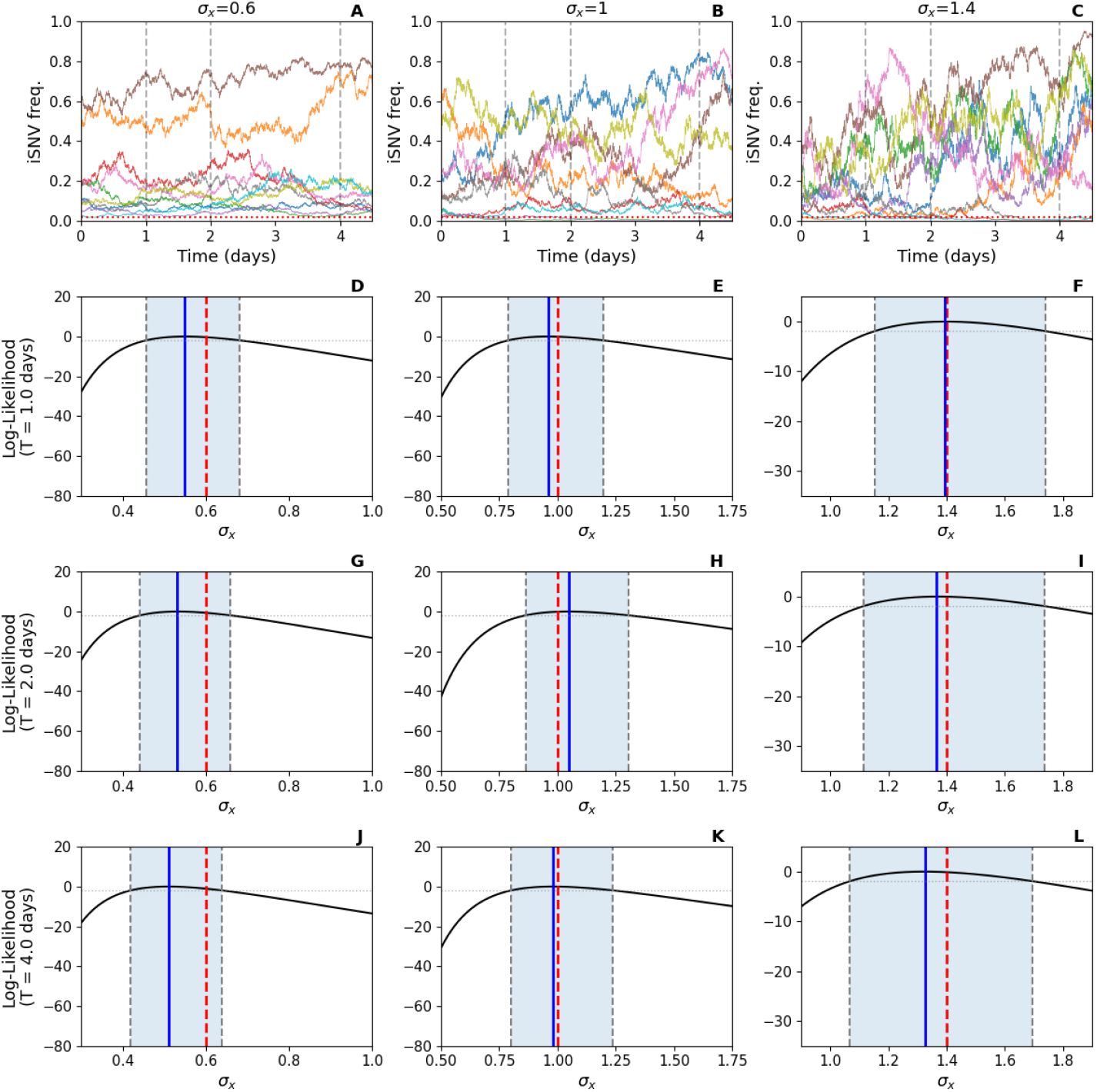
Recovery of *σ*_*x*_ from simulated datasets. (A) A representative set of 10 mock datasets showing simulated iSNV frequency dynamics using *σ*_*x*_ = 0.6. Each dataset shows a simulation of allele frequencies for 4.5 days. Vertical lines are shown at 1 day, 2 days, and 4 days. These are the time points considered for the second collection time. (B) A representative set of 10 mock datasets showing simulated iSNV frequency dynamics using *σ*_*x*_ = 1.0. (C) A representative set of 10 mock datasets showing simulated iSNV frequency dynamics using *σ*_*x*_ = 1.4. (D)-(F) Estimation of *σ*_*x*_ from 50 mock datasets with the second collection time point being *T* = 1 day and the *σ*_*x*_ given by 0.6, 1.0, and 1.4, respectively. (G)-(I) Estimation of *σ*_*x*_ from 50 mock datasets with the second collection time point being *T* = 2 days and the *σ*_*x*_ given by 0.6, 1.0, and 1.4, respectively. (J)-(L) Estimation of *σ*_*x*_ from 50 mock datasets with the second collection time point being *T* = 4 days and the *σ*_*x*_ given by 0.6, 1.0, and 1.4, respectively. In panels (D)-(L), the true *σ*_*x*_ values are shown with dashed red lines, the maximum likelihood estimates of *σ*_*x*_ are shown with solid blue lines, and the 95% CIs are shown using shaded blue regions.

### Estimation of *σ*_*x*_ for acute influenza A virus infections

Figure 4A shows the frequencies of the paired-time IAV iSNV observations listed from Table S1. From these 60 paired-time observations, and assuming that all iSNVs are phenotypically neutral, we estimated *σ*_*x*_ to be *σ*_*x*_ = 1.52 (95% CI = 1.19 - 2.02) (Figure 4B). Downsampling these observations to retain only one paired-time observation per infected individual (to ensure independence of paired-time observation data points) resulted in similar *σ*_*x*_ estimates, although with slightly broader 95% confidence intervals. While we can estimate *σ*_*x*_, the specific value of the extent of environmental noise *ϵ* and the degree of noise correlation *ρ* are not separately identifiable. However, the inferred value of *σ*_*x*_ is consistent with a band of (*ϵ, ρ*) values ranging from relatively low environmental noise *ϵ* and low noise correlation *ρ* to relatively high environmental noise *ϵ* and high noise correlation *ρ* (Figure 4C).

**Figure 4.**
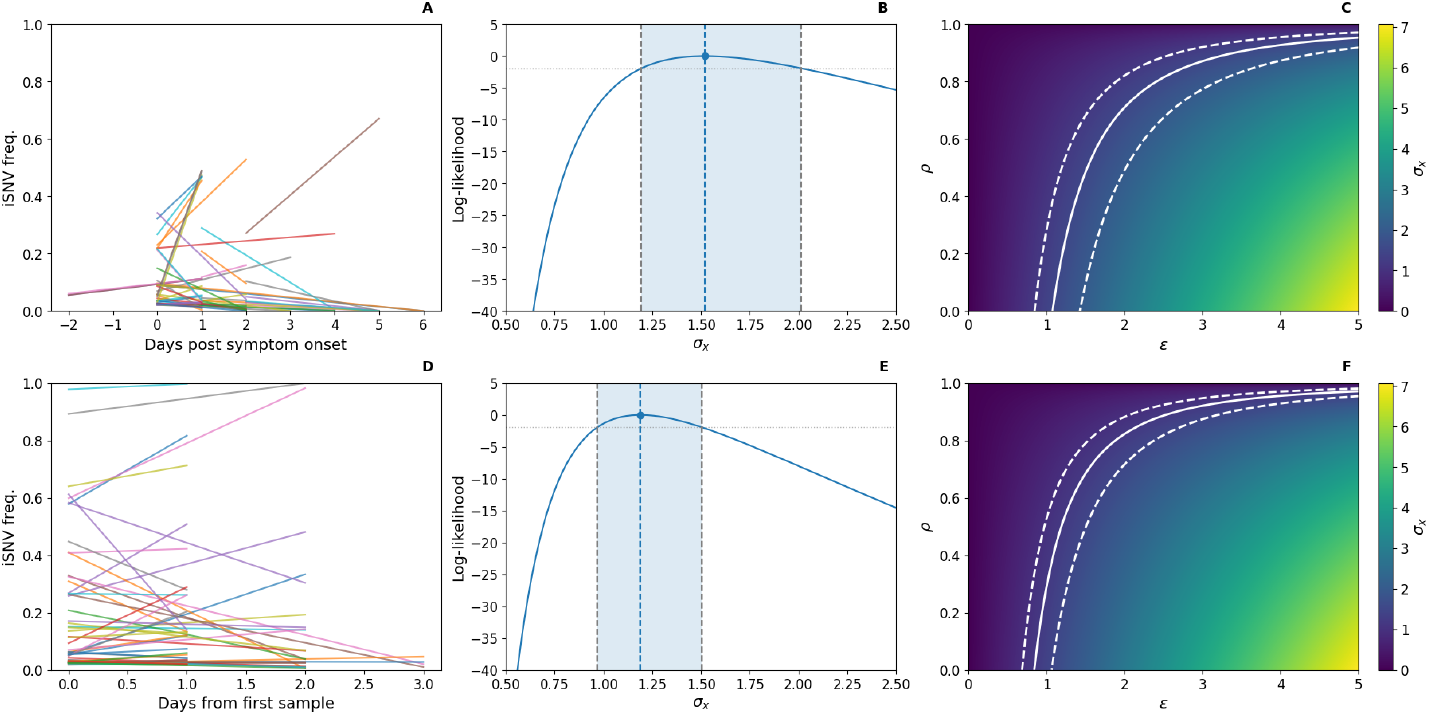
Estimation of *σ*_*x*_ from the influenza A virus data set and from the SARS-CoV-2 data set. (A) Paired-time iSNV observations from the data made publicly available by McCrone et al. (2018). All paired-time observations in Table S1 are plotted. (B) Estimation of *σ*_*x*_ from the paired-time observations shown in panel (A). (C) Parameter combinations of *ϵ* and *ρ* that are consistent with estimates of *σ*_*x*_ for IAV. (D) Paired-time iSNV observations from the data made publicly available by Tonkin-Hill et al. (2021). All paired-time observations in Table S2 are plotted. (E) Estimation of *σ*_*x*_ from the paired-time observations shown in panel (D). (F) Parameter combinations of *ϵ* and *ρ* that are consistent with estimates of *σ*_*x*_ for SARS-CoV-2. In panels (B) and (D), the blue dashed lines show the maximum log-likelihood estimates of *σ*_*x*_ and the 95% confidence intervals are shaded. In panels (C) and (F), parameter combinations of *ϵ* and *ρ* that yield the MLE of *σ*_*x*_ are shown with a solid white line and combinations that fall within the 95% confidence interval of *σ*_*x*_ are shown between the two dashed white lines.

### Estimation of *σ*_*x*_ for acute SARS-CoV-2 infections

Figure 4D shows the frequencies of the paired-time SARS-CoV-2 iSNV observations listed in Table S2. From these 54 paired-time observations, we estimated *σ*_*x*_ to be *σ*_*x*_ = 1.19 (95% CI = 0.97-1.50) (Figure 4E). Downsampling these observations to retain only one paired-time observation per infected individual again resulted in similar *σ*_*x*_ estimates with broader 95% confidence intervals. The specific value of the extent of environmental noise *ϵ* and the degree of noise correlation *ρ* are again not separately identifiable and the inferred value of *σ*_*x*_ is consistent with a band of (*ϵ, ρ*) values ranging from relatively low environmental noise *ϵ* and low noise correlation *ρ* to relatively high environmental noise *ϵ* and high noise correlation *ρ* (Figure 4F). It is interesting to note here the quantitative similarity between the *σ*_*x*_ inferred for IAV and the *σ*_*x*_ inferred for SARS-CoV-2.

### Model criticism using simulated and empirical data

Above, we estimated *σ*_*x*_ from an empirical within-host IAV dataset and an empirical within-host SARS-CoV-2 dataset. We now assess whether our empirical *σ*_*x*_ estimates yield patterns in iSNV frequency changes that are consistent with expected patterns under our evolutionary model that incorporates environmental stochasticity. To do so, we first turn to the mock dataset we simulated to perform model criticism. Figures 5A shows the simulated data, with *X* as the plotted variable. As a reminder, *X* is a quantity that is related to iSNV frequency via equation 7. While the longitudinal simulations of *X* differ between one another, the mean trajectory of *X* remains approximately constant (Figure 5A), as expected, given that 𝔼 [*X*(*t* = *T*)] = *X*(*t* = 0) for our evolutionary model incorporating environmental noise. The spread of *X* trajectories increases with time, reflecting the linear scaling of variance with *T*.

**Figure 5.**
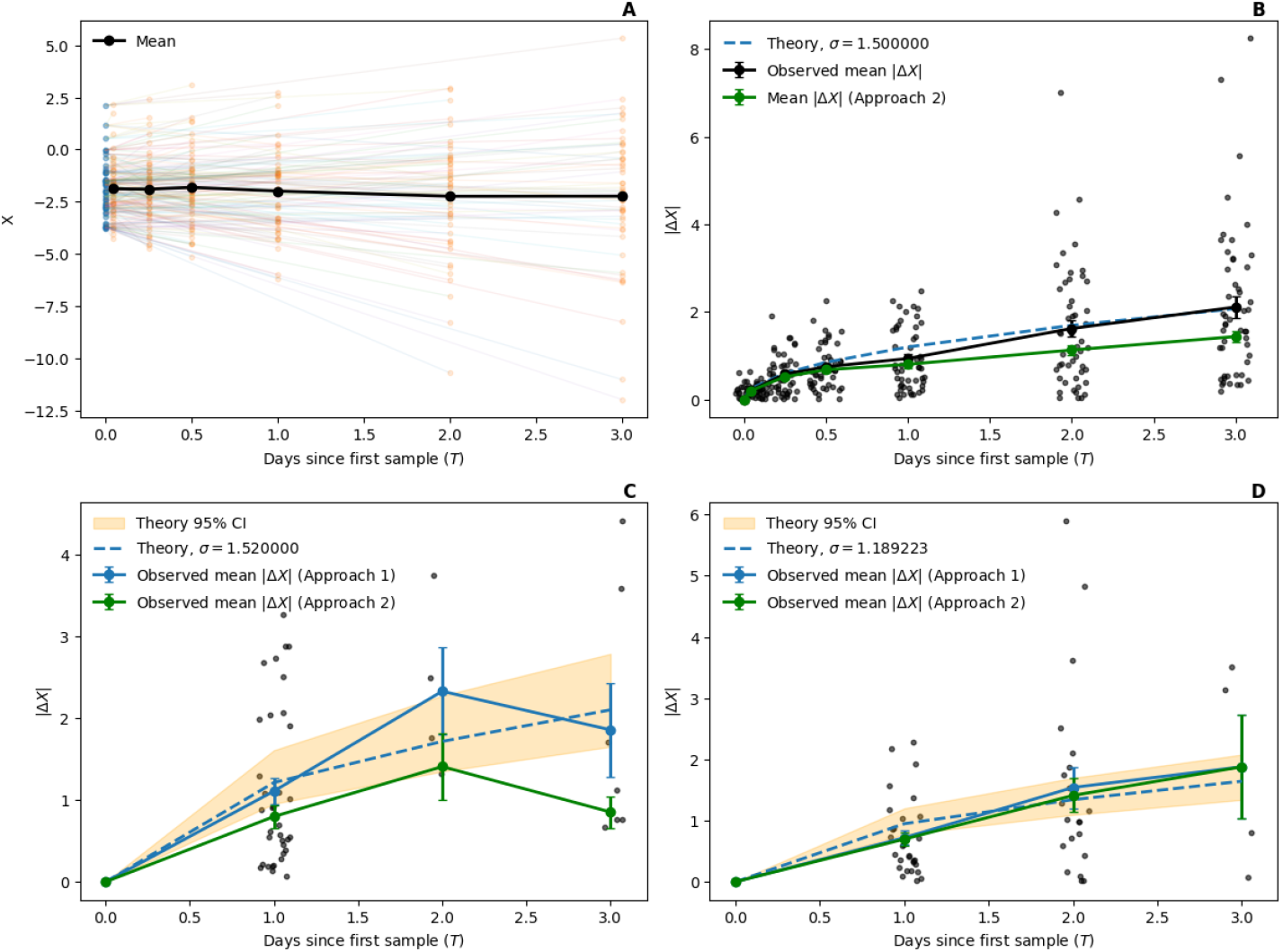
Evaluation of the Brownian motion model using simulated and empirical iSNV frequency data. (A) Dynamics of *X* from the mock dataset to evaluate the Brownian motion model. Here, *σ*_*x*_ was set to 1.5. (B) Absolute changes in *X* from the mock dataset, alongside calculated mean values of these absolute changes and the theoretical prediction of the mean. Black dots show absolute changes in *X* between timepoints from the mock data shown in panel (A). Blue line shows the theoretical mean when *σ*_*x*_ = 1.5, given by equation 10. Black line shows the empirical mean of the blue datapoints. Error bars show the standard error of this mean. Green line shows the empirical mean of the data once second time point frequencies below the variant calling threshold of 2% have been reset to 2%. (C) Absolute changes in *X* from the IAV dataset, alongside calculated mean values of these absolute changes and the theoretical prediction of the mean when *σ*_*x*_ = 1.52. The two paired-time observations with second observed timepoint *T* = 4 days and the two paired-time observation with second observed timepoint *T* = 6 days are excluded from the plot. Δ*X* values for the *T* = 4 day paired-time observations are 0.27 and 2.14. Both *T* = 6 second time point frequencies were 0 and as such there Δ*X* values are not plotted. (D) Absolute changes in *X* from the SARS-CoV-2 dataset, alongside calculated mean values of these absolute changes and the theoretical prediction of the mean when *σ*_*x*_ = 1.19. Δ*X* values from all data points in Table S2 are shown. In panels (C) and (D), the blue lines show mean values when paired-time observations having an iSNV frequency of 0$ at the second time point are excluded (Approach 1). The green lines show mean values when paired-time observations having iSNV frequencies below the variant-calling threshold of 2% being reset at 2% (Approach 2). Error bars on the blue and green lines show standard errors of the means. Dashed blue lines show the theoretical prediction when parameterized with the MLE of *σ*_*x*_ and the shaded orange regions shows theoretical predictions of the mean when considering the 95% CI of *σ*_*x*_. Black dots show Δ*X* values from individual paired-time observations for those observations with *f*_2_(*t* = *T*) *>* 0.

Using this simulated dataset, we then look at whether the observed absolute changes in *X* follows the theoretical prediction. Specifically, we plot in Figure 5B the empirical mean absolute change in *X* as a function of time from the first sample taken at time *t* = 0. This mean closely matches the theoretically expected relationship under the model (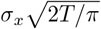 ; Figure 5B). Standard errors of the mean are small due to the large number of observations (50 at each time point). These results indicate that the theoretical expectation provides an accurate description of the dynamics. We further plot the mean after applying Approach 2 to below-the-variant-calling threshold iSNV frequencies taken at the second time point. (That is, resetting all sample 2 iSNV frequencies that fall below 2% to be at 2%.) This results in a mean absolute change in *X* that is slightly lower than the theoretical mean, as expected, given that this resetting has the effect of decreasing some Δ*X* values. We do not apply Approach 1 because none of our simulated iSNV frequencies have *f*_2_(*t* = *T*) = 0.

We next applied the same analysis to our empirical datasets (Figures 5C, D). When excluding paired-time observations that had a second observation frequency of 0% (*n* = 12 for IAV and *n* = 0 for SARS-CoV-2; Approach 1), empirical mean absolute changes in *X* increased with sampling interval *T* in a manner that was highly consistent with that expected from our evolutionary model parameterized with *σ*_*x*_ values of 1.52 and 1.19, respectively. When retaining all paired-time observations but setting the second observation to 2% if the iSNV frequency fell below 2% (Approach 2), empirical mean absolute changes in *X* again increased with sampling interval *T* in a manner that was highly consistent with that expected from our evolutionary model. As expected, the mean values under Approach 2 were slightly lower than those under Approach 1.

Finally, we put both our simulated and empirical data sets to a more stringent test by examining whether the observed distribution of changes in *X* in these data sets were quantitatively similar to those expected theoretically. Figure 6A shows the empirical distribution of changes in *X* from our *T* = 1 day apart paired-time simulated dataset alongside the theoretical prediction, given by a normal distribution with mean 0 and variance 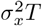. As expected, the empirical distribution appears similar to its theoretical prediction. In Figure 6B, we show both the empirical cumulative distribution function (cdf) and the cdf of the reference normal distribution. A comparison of these cdfs using the one-sample Kolmogorov-Smirnov test yields a p-value of 0.187, indicating no statistical difference between the empirical sample and this reference normal distribution. Figures 6C and 6D perform similar analyses using our *T* = 2 days apart paired-time simulated dataset. Again, we find no statistical difference between the empirical sample and this reference normal distribution. In Supplemental Figure S1, we provide similar analyses for paired time point observations taken 1 hour apart, 6 hours apart, 12 hours apart, and 3 days apart. In all of these cases, and as expected, there is no statistical difference between the empirical sample and the reference normal distributions. We then apply the same analyses to the IAV and SARS-CoV-2 datasets. For IAV, we similarly find no statistical difference between the empirical sample from *T* = 1 day apart observations and this sample’s reference normal distribution (mean 0 and variance calculated with *σ*_*x*_ = 1.52 and *T* = 1 day (Figure 6E, F). Supplemental Figure S2 shows similar analyses for sample sets from *T* = 2 and *T* = 3 day apart observations. (We include these in the supplement because the number of paired-time observations with *T* = 2 and *T* = 3 are very small (*n* = 4 and *n* = 7, respectively). In all of these analyses, we find no statistical difference between the empirical IAV samples and their normal distributions. Turning to SARS-CoV-2, for our *T* = 1 time-paired observations, the K-S test yields a marginally significant p-value (*p* = 0.047) when the sample’s cdf is compared against a normal distribution with mean 0 and variance calculated with *σ*_*x*_ = 1.19 and *T* = 1 (Figure 6G, H). When considering for *T* = 2 time-paired observations, however, the K-S test again yields no statistical differences between this sample and its normal distribution (p-value = 0.265; Figure 6I,J). Supplemental Figure S3 shows similar analyses for the set of paired-time observations that were sampled *T* = 3 days apart (*n* = 4). In this analysis, again there is no statistical difference between the sample and its normal distribution. Overall, these results show that our Brownian motion model, incorporating environmental stochasticity in driving iSNV frequency changes, yields quantitative predictions that are largely consistent with empirical IAV and SARS-CoV-2 within-host data.

**Figure 6.**
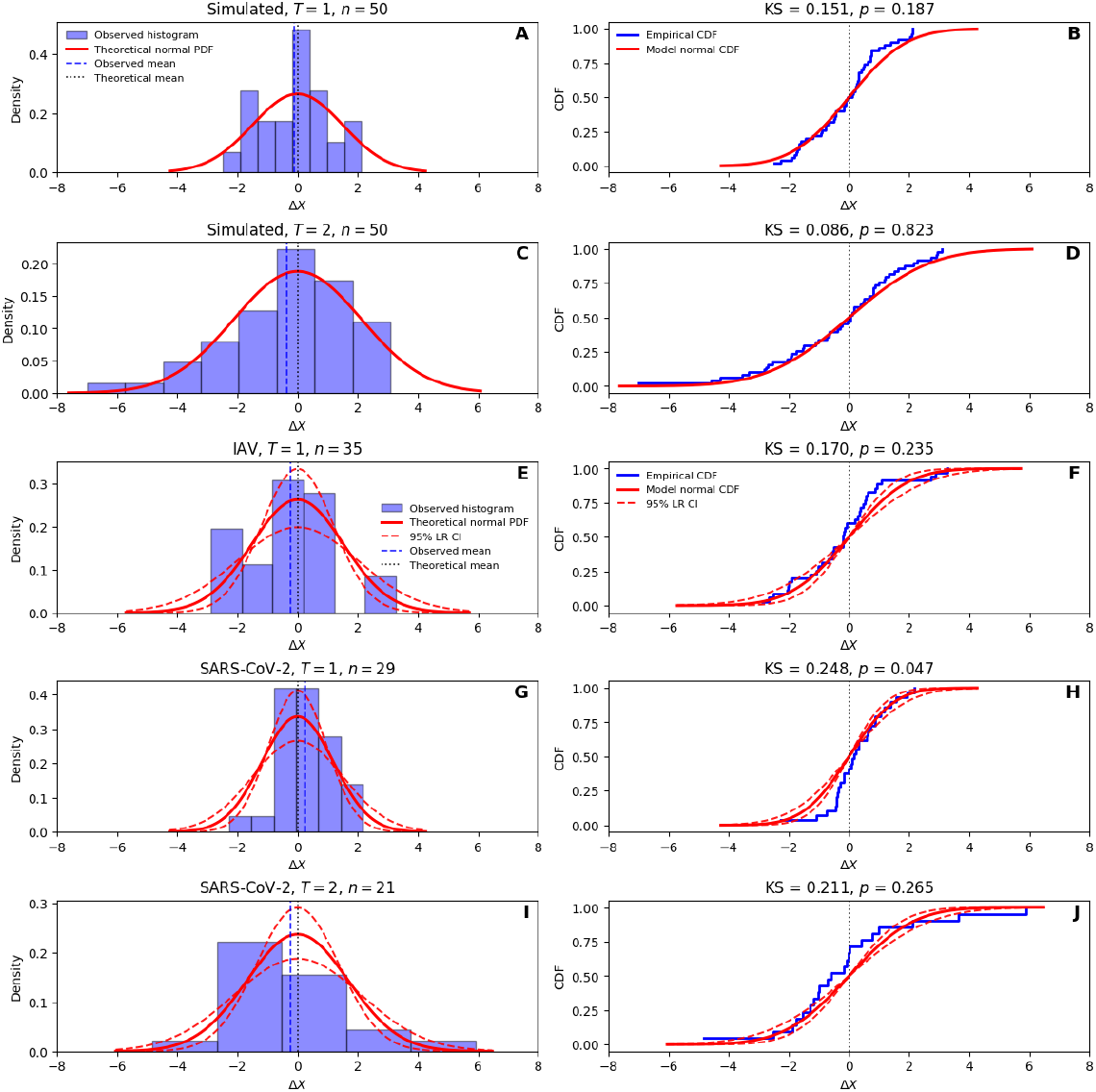
Comparison of observed changes in *X* against theoretical reference distributions. (A) Distribution of simulated changes in *X* between samples taken one day apart (*T* = 1 day) alongside the theoretical reference distribution, given by a normal distribution with mean 0 and variance 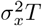. (B) Empirical cumulative distribution function (cdf) for changes in *X* from the empirical dataset shown in panel (A), alongside the cdf of the normal distribution shown in panel (A). Comparison of these cdfs using the one-sample Kolmorogov-Smirnov test yielded a p-value of 0.187. (C) Distribution of simulated changes in *X* between samples taken two days apart (*T* = 2 days). (D) Empirical cdf for changes in *X* from the empirical dataset shown in panel (C), alongside the cdf of the normal distribution shown in panel (C). Comparison of these cdfs using the one-sample Kolmorogov-Smirnov test yielded a p-value of 0.823. (E) Distribution of observed changes in *X* between IAV samples taken two days apart (*T* = 1 day) alongside its theoretical reference distribution, given by a normal distribution with mean 0 and variance 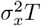, with *σ*_*x*_ = 1.521. (F) Empirical IAV cdf for changes in *X* from the empirical dataset shown in panel (E), alongside the cdf of the normal distribution shown in panel (E). Comparison of these cdfs using the one-sample Kolmorogov-Smirnov test yielded a p-value of 0.235. (G) Distribution of observed changes in *X* between SARS-CoV-2 samples taken one day apart (*T* = 1 day) alongside its theoretical reference distribution, given by a normal distribution with mean 0 and variance 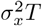, with *σ*_*x*_ = 1.189. (H) Empirical SARS-CoV-2 cdf for changes in *X* from the empirical dataset shown in panel (G), alongside the cdf of the normal distribution shown in panel (G). Comparison of these cdfs using the one-sample Kolmorogov-Smirnov test yielded a p-value of 0.047. (I) Distribution of observed changes in *X* between SARS-CoV-2 samples taken two days apart (*T* = 2 days) alongside its theoretical reference distribution, given by a normal distribution with mean 0 and variance 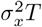, with *σ*_*x*_ = 1.189. (J) Empirical SARS-CoV-2 cdf for changes in *X* from the empirical dataset shown in panel (I), alongside the cdf of the normal distribution shown in panel (I). Comparison of these cdfs using the one-sample Kolmorogov-Smirnov test yielded a p-value of 0.265.

### Selection remains possible within acutely infected individuals in the presence of environmental stochasticity

Here, we examine frequency dynamics of iSNVs that are not fitness neutral. Figure 7A shows simulations of the Arithmetic Brownian Motion model given by equation 12 when the focal iSNV has a considerable selective advantage (*r*_*x*_ = 1). Three sets of simulations are shown: one with *σ*_*x*_ = 0.6, one with *σ*_*x*_ = 1.0, and one with *σ*_*x*_ = 1.4, all with the focal iSNV initially present at time *t* = 0 at a frequency of 1%. From this plot, it is clear that an iSNV with a selective advantage of this magnitude can increase in frequency over a three-day time period. Despite the iSNV’s selective advantage, not all simulations result in the detection of the iSNV above the 2% variant calling threshold. Indeed, the higher the amount of environmental noise, the lower the probability that the iSNV will be observed above the 2% threshold by time *T* = 3 (Figure 7C). These results indicate that positive selection can act on standing genetic variation in the presence of biologically relevant amounts of stochasticity. This finding is consistent with experimental challenge studies that have shown evidence of within-host viral positive selection (Sobel Leonard et al., 2016; Ferreri et al., 2026; Wilker et al., 2013; Plante et al., 2021; Zhou et al., 2021). The lack of detection of positive selection in natural infections (McCrone et al., 2018; Dinis et al., 2016) may thus indicate that beneficial variants are too late in arising or too small in selective advantage to be able to have selection bring them to appreciable levels over the time course of an acute infection.

**Figure 7.**
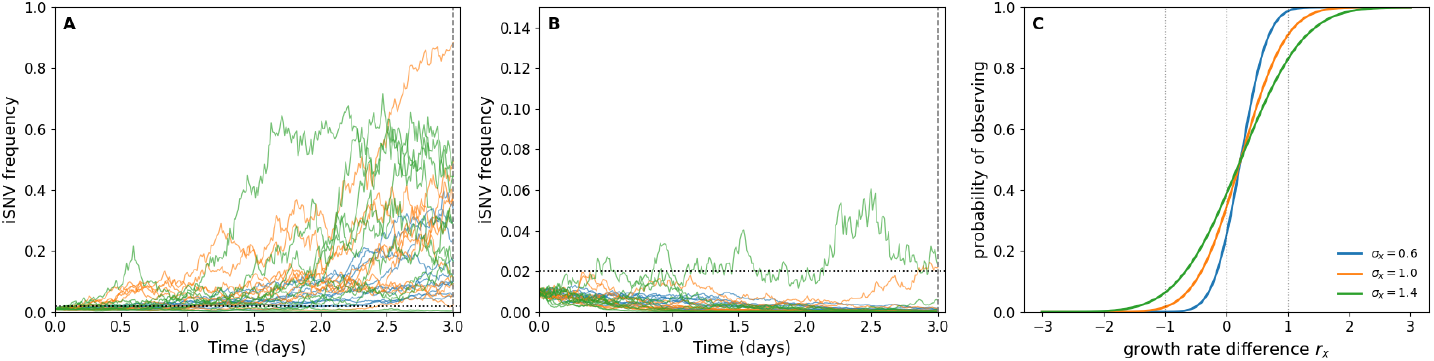
Simulated dynamics of beneficial and deleterious iSNVs. (A) Simulations of beneficial iSNVs with *r*_*x*_ = 1. (B) Simulations of deleterious iSNVs with *r*_*x*_ = −1. (A, B) Three sets of 10 simulated iSNV frequency dynamics are shown: *σ*_*x*_ = 0.6 (blue) *σ*_*x*_ = 1 (orange), and *σ*_*x*_ = 1.4 (green). Initial iSNV frequencies are set to 0.01. (C) Analytical probability of observing an iSNV at sampling time *T* = 3 days as a function of *r*_*x*_, the growth rate difference between the focal iSNV and the alternative allele. An iSNV is considered observed if its frequency at time *T* is above the variant-calling threshold of 2%. Vertical lines are shown at *r*_*x*_ = -1, 0, and 1. *r*_*x*_ = 1 and *r*_*x*_ = −1 are the values used in panels (A) and (B), respectively.

Figure 7B shows simulations of the Arithmetic Brownian Motion model given by equation 12 when the focal iSNV has a selective disadvantage (*r*_*x*_ = −1). Three sets of simulations are again shown at values of *σ*_*x*_ = 0.6, 1.0, and 1.4, with the focal iSNV initially present at time *t* = 0 at a frequency of 1%. From this plot, it is clear that an iSNV with a selective disadvantage of this magnitude usually decreases in frequency over a three-day time period. Again, when *σ*_*x*_ = 0.6, the dynamics of the iSNV appear most deterministic, as expected. Despite the iSNV’s selective disadvantage, not all simulations result in the iSNV staying below the 2% variant calling threshold. Indeed, the higher the amount of environmental noise, the higher the probability that the iSNV will be observed above the 2% threshold by time *T* = 3 (Figure 7C). These results indicate that purifying selection can act on standing genetic variation in the presence of considerable amounts of stochasticity. This finding is consistent with findings from natural infections (McCrone et al., 2018; Lythgoe et al., 2021).

## Discussion

The evolution of respiratory viruses within acute human infections has previously been described as highly stochastic, with large changes in allele frequencies possible within a matter of days (McCrone et al., 2018; Tonkin-Hill et al., 2021; Lythgoe et al., 2021). To quantify the magnitude of this stochasticity, Wright-Fisher type models have been applied to within-host viral data to estimate within-host viral effective population sizes (McCrone et al., 2020; Lumby et al., 2020; Shi et al., 2026). However, while it is clear that these models can capture and reproduce key quantitative features of observed within-host viral dynamics, it is unclear how to interpret a single estimate of *N*_*E*_ for within-host infections, where the census viral population size first increases and then decreases by many orders of magnitude. Moreover, genetic drift in these Wright-Fisher models is assumed to stem from demographic stochasticity. Such demographic stochasticity is expected to become less pronounced in larger populations. Given the extremely large viral population sizes at and around the peak of infection, it becomes difficult to believe that demographic stochasticity is the primary driver of allele frequency changes during these time periods. Here, we therefore considered environmental stochasticity as the non-selective process driving within-host viral evolution. By developing and analyzing models incorporating environmental noise, we showed how allele frequencies could change rapidly when the magnitude of environmental noise *ϵ* was sufficiently large, provided that the correlation of noise between circulating strains *ρ* is not perfect (*ρ* ≠ 1). We further developed a statistical method to interface with allele frequency data to infer a quantity *σ*_*x*_. Technically speaking, this parameter captures the magnitude of Brownian motion diffusion and reflects a combination of *ϵ* and *ρ*. We applied this method to empirical IAV and SARS-CoV-2 datasets, estimating *σ*_*x*_ and further showing that their observed evolutionary dynamics were consistent with our model incorporating environmental noise. Finally, we showed that both positive and purifying selection would still be able to act in the context of environmental noise.

The models and analyses we developed here focused primarily on understanding the within-host frequency dynamics of neutral alleles in acute respiratory infections. There are several questions we would like to address in future work. These include: Are within-host viral dynamics as noisy during viral establishment and proliferation as they are during viral decline? Most of the within-host longitudinal datasets available from infected individuals derive from individuals who were sampled following symptom onset, which generally corresponds to the viral decline phase. Because of lower viral loads early on in infection, it may be the case that the extent of noise correlation *ρ* is lower during viral proliferation than during the viral clearance phase that follows a high viral peak and potentially therefore more thorough mixing of different viral genotypes. Different processes also dominate during viral proliferation versus viral decline. For example, if environmental noise related to viral infection of susceptible target cells is large, then this noise would have the opportunity to act during viral proliferation. However, during viral decline, noise related to this process would not impact viral dynamics because viral infection no longer plays a large role in regulating viral population sizes. Because the key processes driving viral within-host dynamics change throughout the course of an acute infection, and the extent and correlation of noise can differ by process, allele frequency dynamics may differ between the viral proliferation period and the viral decline period. By fitting our Brownian motion model to data from these two different infection ‘epochs’, we could start to gauge whether allele frequency dynamics quantitatively differ from one another during these periods.

Another set of questions that our model could address is how environmental noise shapes allele frequency dynamics in chronic or prolonged respiratory virus infections. In many documented instances of these infections, positive selection has been detected, with iSNVs conferring either immune escape or other fitness-enhancing traits observed to rise in frequency (Kemp et al., 2021; Scherer et al., 2022). In cases such as these, we could use our expanded model with selection and determine how in these infections natural selection and changes in allele frequencies driven by environmental stochasticity jointly impact patterns of viral evolution. This approach would also be possible in acute infections from experimental challenge studies where selection and drift have both been apparent (Sobel Leonard et al., 2016; Ferreri et al., 2026). Understanding of other viral infections, such as enteric viruses (e.g., noroviruses) and systemically circulating viruses (e.g., HIV and HCV), may also benefit from application of this environmental stochasticity model.

Our current approach is limited to paired-time observations of a single iSNV within the same individual. While this limitation is not overly restrictive in application to natural, low-diversity viral infections, it is limiting in the case of viral populations with considerably more within-host diversity This includes barcoded viral populations that have increasingly been used to study within-host and between-host viral evolutionary dynamics (Varble et al., 2014; Amato et al., 2022; Holmes et al., 2025; Ferreri et al., 2025). In this case, instead of two strains, there are hundreds to thousands of barcoded viruses, each of which we can consider a strain. We are currently extending our model to consider viral populations such as these. Intriguingly, several existing studies have shown the maintenance of viral barcode diversity throughout the course of infection in inoculated animals (e.g., Holmes et al. (2025); Ferreri et al. (2025)), with barcode frequencies that sometimes only change very little between timepoints. Dynamics such as these could be considered more quantitatively with models such as the one presented here, providing insight into why evolutionary differences might exist between experimentally challenged animals and natural infections. For example, barcoded viruses in the viral inoculum of these studies are well-mixed and a large inoculum volume (with a mode of delivery that involves instillation of bulk liquid) might seed these barcodes throughout the respiratory tract, with single barcodes distributed at random across the areas where viruses deposited. Even though viruses in different spatial locations might still experience environmental noise to the same degree as in a natural infection, noise in a specific location would be felt by many different barcodes. As such, averaging across space, individual viral barcodes may effectively experience noise in a manner that is highly similar to other barcodes. This would result in an overall high level of noise correlation (*ρ* close to 1) between barcoded viruses and thus the expectation that observed barcode frequencies would not change appreciably.

Finally, our derived model assumes that environmental noise is white noise, that is, uncorrelated noise. In the ecological literature, noise is known to come in different ‘colors’. While the color of noise in terrestrial systems appears to be white, the color of noise in marine systems appears to be red or Brown (Steele, 1985; Vasseur and Yodzis, 2004). Red noise is strong in longer wavelengths and Brown (or Brownian) noise refers to random walk noise. Through extensions of the types of models presented here, we could examine whether the respiratory tract environment subjects viral populations not only to noise but to a certain color of noise. Answering these questions will require sufficiently detailed longitudinal within-host data sets and collaboration between experimentalists and modelers to effectively query these data for improved understanding of the processes driving within-host viral evolution and adaptation.

## Appendix

### Derivation of the Brownian motion model

Taking the derivative of *X* in equation 6 yields *dX*(*t*) = *d* log *V*_2_(*t*) − *d* log *V*_1_(*t*). We have expressions for *d* log *V*_1_(*t*) and *d* log *V*_2_(*t*) by log-transforming equations 3 and 4 using Itô calculus:

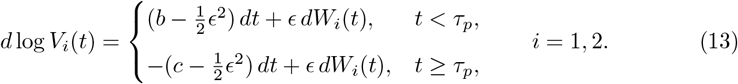

Substituting into the equation for *dX*(*t*) yields:

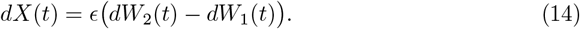

The expected value of the difference (*dW*_2_(*t*)−*dW*_1_(*t*)) is 0. The variance of the difference (*dW*_2_(*t*) − *dW*_1_(*t*)) can be calculated using the identity *V ar*(*A* − *B*) = *V ar*(*A*)+*V ar*(*B*) − 2*Cov*(*A, B*):

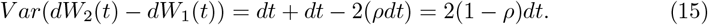

We can therefore express equation 18 using a Brownian motion model of the form *dX*(*t*) = *σ*_*x*_*dW*_*x*_(*t*) where *dW*_*x*_(*t*) is white noise, drawn from a normal distribution with mean 0 and variance *dt*. Parameter *σ*_*x*_ quantifies the intensity of the random fluctuations and can be written as a function of *ρ* and parameter *ϵ*. To do so, we first write the difference (*dW*_2_(*t*) − *dW*_1_(*t*)) as a scaled standard Brownian increment:

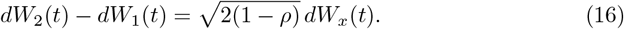

This preserves the variance given in equation 15. Substituting equation (16) into equation (18) yields 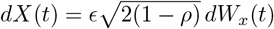. As such, the dynamics of variable *X*(*t*) follow a Brownian motion model with diffusion parameter (equation 9).

#### Derivation of the non-neutral evolutionary model

In the piecewise exponential, strains 1 and 2 are non-neutral if their proliferation rates (*b* values) or if their clearance rates (*c* values) differ from one another. We can thus make both *b* and *c* a function of strain (*b*_*i*_ and *c*_*i*_). If *b*_2_ *> b*_1_, then strain 2 has a selective advantage during the viral proliferation period. If *b*_2_ *< b*_1_, then strain 2 has a selective disadvantage during the viral proliferation period. Similarly, if *c*_2_ *< c*_1_, then strain 2 has a selective advantage during the viral decline period. If *c*_2_ *> c*_1_, then strain 2 has a selective disadvantage during the viral decline period. Substituting into the equation *dX*(*t*) = *d* log *V*_2_(*t*) − *d* log *V*_1_(*t*) yields:

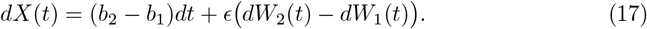

during the expansion phase (*t < τ*_*p*_) and

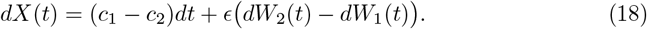

during the decline phase (*t > τ*_*p*_). Simplifying the second terms on the right hand side according to our analyses in the above subsection yields:

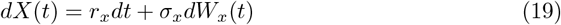

where *r*_*x*_ is defined as either *b*_2_ − *b*_1_ (if *t < τ*_*p*_) or *c*_1_ − *c*_2_ (if *t > τ*_*p*_). This model is an Arithmetic Brownian Motion model with drift term *r*_*x*_*dt* and diffusion term *σ*_*x*_*dW*_*x*_(*t*).

## Supplemental figure legends

**Supplementary Figure S1.**
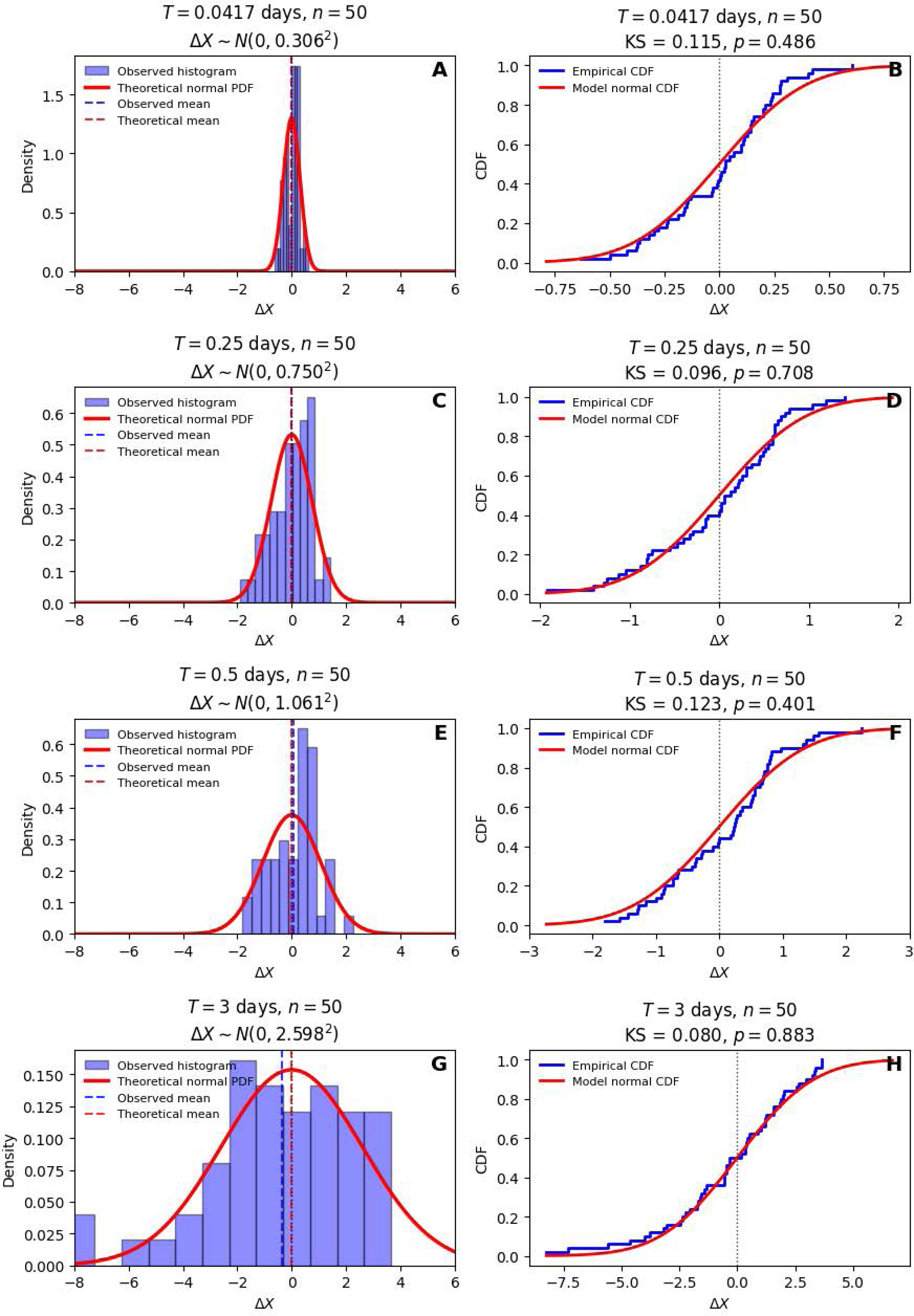
Comparison of observed changes in *X* from the simulated dataset against theoretical reference distributions. (A) Distribution of simulated changes in *X* between samples taken 1 hour apart (*T* = 0.0417 days) alongside the theoretical reference distribution, given by a normal distribution with mean 0 and variance 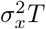. (B) Empirical cumulative distribution function (cdf) for changes in *X* from the empirical dataset shown in panel (A), alongside the cdf of the normal distribution shown in panel (A). Comparison of these cdfs using the one-sample Kolmorogov-Smirnov test yielded a p-value of 0.486. (C) Distribution of simulated changes in *X* between samples taken 6 horus apart (*T* = 0.25 days). (D) Empirical cdf for changes in *X* from the empirical dataset shown in panel (C), alongside the cdf of the normal distribution shown in panel (C). Comparison of these cdfs using the one-sample Kolmorogov-Smirnov test yielded a p-value of 0.708. (E) Distribution of simulated changes in *X* between samples taken half a day apart (*T* = 0.5 days). (F) Empirical cdf for changes in *X* from the empirical dataset shown in panel (E), alongside the cdf of the normal distribution shown in panel (E). Comparison of these cdfs using the one-sample Kolmorogov-Smirnov test yielded a p-value of 0.401. (G) Distribution of simulated changes in *X* between samples taken three days apart (*T* = 3 days). (H) Empirical cdf for changes in *X* from the empirical dataset shown in panel (G), alongside the cdf of the normal distribution shown in panel (G). Comparison of these cdfs using the one-sample Kolmorogov-Smirnov test yielded a p-value of 0.883.

**Supplementary Figure S2.**
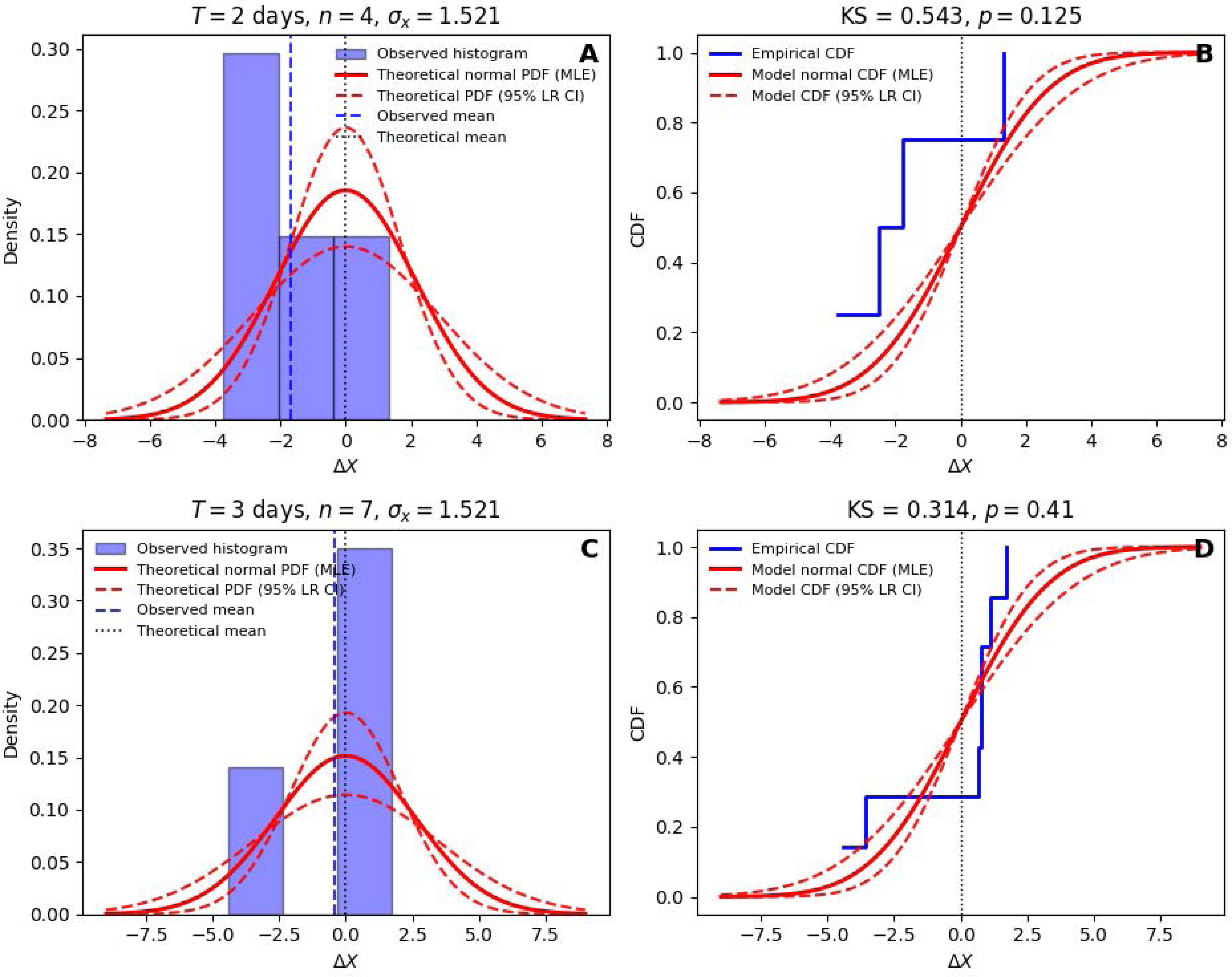
Comparison of observed changes in *X* in the IAV dataset against theoretical reference distributions. (A) Distribution of observed changes in *X* between samples taken two days apart (*T* = 2 days). (B) Empirical cdf for changes in *X* from the empirical dataset shown in panel (A), alongside the cdf of the normal distribution shown in panel (A). Comparison of these cdfs using the one-sample Kolmorogov-Smirnov test yielded a p-value of 0.125. (C) Distribution of observed changes in *X* between samples taken three days apart (*T* = 3 days). (D) Empirical cdf for changes in *X* from the empirical dataset shown in panel (C), alongside the cdf of the normal distribution shown in panel (C). Comparison of these cdfs using the one-sample Kolmorogov-Smirnov test yielded a p-value of 0.41.

**Supplementary Figure S3.**
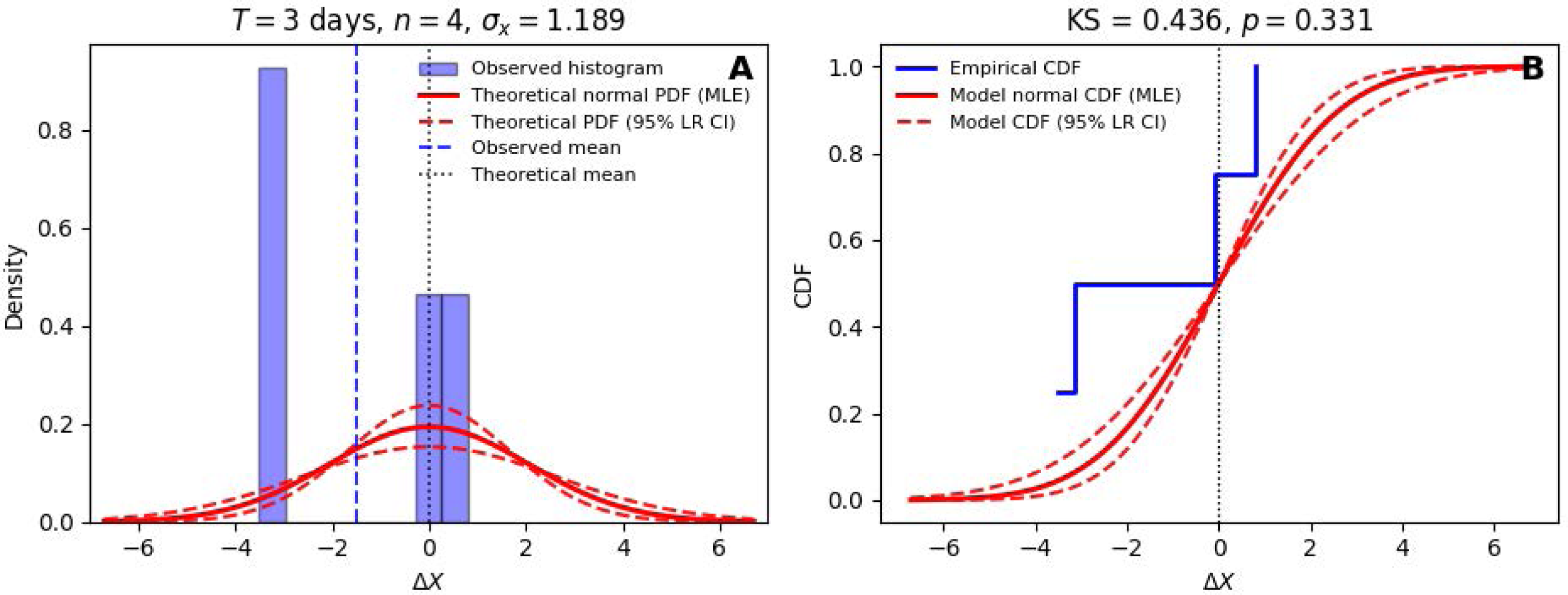
Comparison of observed changes in *X* in the SARS-CoV-2 dataset against theoretical reference distributions. (A) Distribution of observed changes in *X* between samples taken three days apart (*T* = 3 days). (B) Empirical cdf for changes in *X* from the empirical dataset shown in panel (A), alongside the cdf of the normal distribution shown in panel (A). Comparison of these cdfs using the one-sample Kolmorogov-Smirnov test yielded a p-value of 0.331.

## References

Alnaji, F. G. and Brooke, C. B. 2020. Influenza virus DI particles: Defective interfering or delightfully interesting? PLoS Pathog., 16(5): e1008436.

Amato, K. A., Haddock, 3rd, L. A., Braun, K. M., Meliopoulos, V., Livingston, B., Honce, R., Schaack, G. A., Boehm, E., Higgins, C. A., Barry, G. L., Koelle, K., Schultz-Cherry, S., Friedrich, T. C., and Mehle, A. 2022. Influenza a virus undergoes compartmentalized replication in vivo dominated by stochastic bottlenecks. Nat. Commun., 13(1): 3416.

Baccam, P., Beauchemin, C., Macken, C. A., Hayden, F. G., and Perelson, A. S. 2006. Kinetics of influenza a virus infection in humans. J. Virol., 80(15): 7590–7599.

Bisht, K. and Te Velthuis, A. J. W. 2022. Decoding the role of temperature in RNA virus infections. MBio, 13(5): e0202122.

Brooke, C. B., Ince, W. L., Wrammert, J., Ahmed, R., Wilson, P. C., Bennink, J. R., and Yewdell, J. W. 2013. Most influenza a virions fail to express at least one essential viral protein. J. Virol., 87(6): 3155–3162.

Coulson, T., Rohani, P., and Pascual, M. 2004. Skeletons, noise and population growth: the end of an old debate? Trends Ecol. Evol., 19(7): 359–364.

Diesel, D. A., Lebel, J. L., and Tucker, A. 1991. Pulmonary particle deposition and airway mucociliary clearance in cold-exposed calves. Am. J. Vet. Res., 52(10): 1665–1671.

Dinis, J. M., Florek, K. R., Fatola, O. O., Moncla, L. H., Mutschler, J. P., Charlier, O. K., Meece, J. K., Belongia, E. A., and Friedrich, T. C. 2016. Deep sequencing reveals potential antigenic variants at low frequencies in influenza a virus-infected humans. J. Virol., 90(7): 3355–3365.

Evans, S. S., Repasky, E. A., and Fisher, D. T. 2015. Fever and the thermal regulation of immunity: the immune system feels the heat. Nat. Rev. Immunol., 15(6): 335–349.

Ferreri, L. M., Seibert, B., Caceres, C. J., Patatanian, K., Holmes, K. E., Gay, L. C., Cargnin Faccin, F., Cardenas, M., Carnaccini, S., Shetty, N., Rajao, D., Koelle, K., Marr, L. C., Perez, D. R., and Lowen, A. C. 2025. Dispersal of influenza virus populations within the respiratory tract shapes their evolutionary potential. Proc. Natl. Acad. Sci. U. S. A., 122(4): e2419985122.

Ferreri, L. M., Vargas-Maldonado, N., Le Sage, V., Leyson, C. M., Pauly, M. D., VanInsberghe, D., Raghunathan, V., Danzy, S., Macenczak, H., Traenkner, J., Flu CHIM Study Group, Catchpole, A., Mann, A., Rouphael, N. G., Lakdawala, S. S., Koelle, K., and Lowen, A. C. 2026. Within-host adaptive evolution is limited by genetic drift in experimental human influenza a virus infections.

Fortier, E. E., Rooney, J., Dardente, H., Hardy, M.-P., Labrecque, N., and Cermakian, N. 2011. Circadian variation of the response of T cells to antigen. J. Immunol., 187(12): 6291–6300.

Foxman, E. F., Storer, J. A., Fitzgerald, M. E., Wasik, B. R., Hou, L., Zhao, H., Turner, P. E., Pyle, A. M., and Iwasaki, A. 2015. Temperature-dependent innate defense against the common cold virus limits viral replication at warm temperature in mouse airway cells. Proc. Natl. Acad. Sci. U. S. A., 112(3): 827–832.

Fujiwara, M. and Takada, T. 2017. Environmental Stochasticity, pages 1–8. John Wiley & Sons, Ltd, Chichester, UK.

Galani, I. E., Triantafyllia, V., Eleminiadou, E.-E., Koltsida, O., Stavropoulos, A., Manioudaki, M., Thanos, D., Doyle, S. E., Kotenko, S. V., Thanopoulou, K., and Andreakos, E. 2017. Interferon-λ mediates non-redundant front-line antiviral protection against influenza virus infection without compromising host fitness. Immunity, 46(5): 875–890.e6.

Gillespie, D. T. 1977. Exact stochastic simulation of coupled chemical reactions. J. Phys. Chem., 81(25): 2340–2361.

Gillespie, D. T. 2001. Approximate accelerated stochastic simulation of chemically reacting systems. J. Chem. Phys., 115(4): 1716–1733.

Gong, H. H., Worley, M. J., Carver, K. A., Godin, C. J., and Deng, J. C. 2025. Deficient neutrophil responses early in influenza infection promote viral replication and pulmonary inflammation. PLoS Pathog., 21(1): e1012449.

Hagan, T., Cortese, M., Rouphael, N., Boudreau, C., Linde, C., Maddur, M. S., Das, J., Wang, H., Guthmiller, J., Zheng, N.-Y., Huang, M., Uphadhyay, A. A., Gardinassi, L., Petitdemange, C., McCullough, M. P., Johnson, S. J., Gill, K., Cervasi, B., Zou, J., Bretin, A., Hahn, M., Gewirtz, A. T., Bosinger, S. E., Wilson, P. C., Li, S., Alter, G., Khurana, S., Golding, H., and Pulendran, B. 2019. Antibiotics-driven gut microbiome perturbation alters immunity to vaccines in humans. Cell, 178(6): 1313–1328.e13.

Hay, J. A., Laurie, K., White, M., and Riley, S. 2019. Characterising antibody kinetics from multiple influenza infection and vaccination events in ferrets. PLoS Comput. Biol., 15(8): e1007294.

Holmes, K. E., Ferreri, L. M., Elie, B., Ganti, K., Lee, C.-Y., VanInsberghe, D., and Lowen, A. C. 2025. Viral expansion after transfer is a primary driver of influenza a virus transmission bottlenecks. PLoS Biol., 23(9): e3003352.

Jacobs, N. T., Onuoha, N. O., Antia, A., Steel, J., Antia, R., and Lowen, A. C. 2019. Incomplete influenza a virus genomes occur frequently but are readily complemented during localized viral spread. Nat. Commun., 10(1): 3526.

Jiang, S., Freedman, J., Mantri, M., Maymi, V., Leddon, S. A., Schweitzer, P., Bhandari, S., Holdener, C., Ntekas, I., Vollmers, C., Flyak, A. I., Fowell, D. J., Rudd, B. D., and De Vlaminck, I. 2026. A temporal and spatial atlas of adaptive immune responses in the lymph node following viral infection. Proc. Natl. Acad. Sci. U. S. A., 123(5): e2504742123.

Ke, R., Zitzmann, C., Ho, D. D., Ribeiro, R. M., and Perelson, A. S. 2021. In vivo kinetics of SARS-CoV-2 infection and its relationship with a person’s infectiousness. Proc. Natl. Acad. Sci. U. S. A., 118(49): e2111477118.

Keller, M., Mazuch, J., Abraham, U., Eom, G. D., Herzog, E. D., Volk, H.-D., Kramer, A., and Maier, B. 2009. A circadian clock in macrophages controls inflammatory immune responses. Proc. Natl. Acad. Sci. U. S. A., 106(50): 21407–21412.

Kemp, S. A., Collier, D. A., Datir, R. P., Ferreira, I. A. T. M., Gayed, S., Jahun, A., Hosmillo, M., Rees-Spear, C., Mlcochova, P., Lumb, I. U., Roberts, D. J., Chandra, A., Temperton, N., CITIID-NIHR BioResource COVID-19 Collaboration, COVID-19 Genomics UK (COG-UK) Consortium, Sharrocks, K., Blane, E., Modis, Y., Leigh, K. E., Briggs, J. A. G., van Gils, M. J., Smith, K. G. C., Bradley, J. R., Smith, C., Doffinger, R., Ceron-Gutierrez, L., Barcenas-Morales, G., Pollock, D. D., Goldstein, R. A., Smielewska, A., Skittrall, J. P., Gouliouris, T., Goodfellow, I. G., Gkrania-Klotsas, E., Illingworth, C. J. R., McCoy, L. E., and Gupta, R. K. 2021. SARS-CoV-2 evolution during treatment of chronic infection. Nature, 592(7853): 277–282.

Kissler, S. M., Fauver, J. R., Mack, C., Olesen, S. W., Tai, C., Shiue, K. Y., Kalinich, C. C., Jednak, S., Ott, I. M., Vogels, C. B. F., Wohlgemuth, J., Weisberger, J., DiFiori, J., Anderson, D. J., Mancell, J., Ho, D. D., Grubaugh, N. D., and Grad, Y. H. 2021a. Viral dynamics of acute SARS-CoV-2 infection and applications to diagnostic and public health strategies. PLoS Biol., 19(7): e3001333.

Kissler, S. M., Fauver, J. R., Mack, C., Tai, C. G., Breban, M. I., Watkins, A. E., Samant, R. M., Anderson, D. J., Metti, J., Khullar, G., Baits, R., MacKay, M., Salgado, D., Baker, T., Dudley, J. T., Mason, C. E., Ho, D. D., Grubaugh, N. D., and Grad, Y. H. 2021b. Viral dynamics of SARS-CoV-2 variants in vaccinated and unvaccinated persons. N. Engl. J. Med., 385(26): 2489–2491.

Kloeden, P. E. and Platen, E. 1992a. Numerical solution of stochastic differential equations. Stochastic Modelling and Applied Probability. Springer, Berlin, Germany, 1 edition.

Kloeden, P. E. and Platen, E. 1992b. Numerical solution of stochastic differential equations. Stochastic Modelling and Applied Probability. Springer, Berlin, Germany, 1 edition.

Kudo, E., Song, E., Yockey, L. J., Rakib, T., Wong, P. W., Homer, R. J., and Iwasaki, A. 2019. Low ambient humidity impairs barrier function and innate resistance against influenza infection. Proc. Natl. Acad. Sci. U. S. A., 116(22): 10905–10910.

Lande, R. 1993. Risks of population extinction from demographic and environmental stochasticity and random catastrophes. Am. Nat., 142(6): 911–927.

Laserson, U., Vigneault, F., Gadala-Maria, D., Yaari, G., Uduman, M., Vander Heiden, J. A., Kelton, W., Taek Jung, S., Liu, Y., Laserson, J., Chari, R., Lee, J.-H., Bachelet, I., Hickey, B., Lieberman-Aiden, E., Hanczaruk, B., Simen, B. B., Egholm, M., Koller, D., Georgiou, G., Kleinstein, S. H., and Church, G. M. 2014. High-resolution antibody dynamics of vaccine-induced immune responses. Proc. Natl. Acad. Sci. U. S. A., 111(13): 4928–4933.

Lumby, C. K., Zhao, L., Breuer, J., and Illingworth, Jr, C. 2020. A large effective population size for established within-host influenza virus infection. Elife, 9(e56915): e56915.

Lythgoe, K. A., Hall, M., Ferretti, L., de Cesare, M., MacIntyre-Cockett, G., Trebes, A., Andersson, M., Otecko, N., Wise, E. L., Moore, N., Lynch, J., Kidd, S., Cortes, N., Mori, M., Williams, R., Vernet, G., Justice, A., Green, A., Nicholls, S. M., Ansari, M. A., Abeler-Dörner, L., Moore, C. E., Peto, T. E. A., Eyre, D. W., Shaw, R., Simmonds, P., Buck, D., Todd, J. A., Oxford Virus Sequencing Analysis Group (OVSG), Connor, T. R., Ashraf, S., da Silva Filipe, A., Shepherd, J., Thomson, E. C., COVID-19 Genomics UK (COG-UK) Consortium, Bonsall, D., Fraser, C., and Golubchik, T. 2021. SARS-CoV-2 within-host diversity and transmission. Science, 372(6539): eabg0821.

Mace, T. A., Zhong, L., Kilpatrick, C., Zynda, E., Lee, C.-T., Capitano, M., Minderman, H., and Repasky, E. A. 2011. Differentiation of CD8+ T cells into effector cells is enhanced by physiological range hyperthermia. J. Leukoc. Biol., 90(5): 951–962.

Martin, B. E., Harris, J. D., Sun, J., Koelle, K., and Brooke, C. B. 2020. Cellular co-infection can modulate the efficiency of influenza a virus production and shape the interferon response. PLoS Pathog., 16(10): e1008974.

Martin, M. A. and Koelle, K. 2021. Comment on “genomic epidemiology of superspreading events in austria reveals mutational dynamics and transmission properties of SARS-CoV-2”. Sci. Transl. Med., 13(617): eabh1803.

McCrone, J. T., Woods, R. J., Martin, E. T., Malosh, R. E., Monto, A. S., and Lauring, S. 2018. Stochastic processes constrain the within and between host evolution of influenza virus. Elife, 7(e35962).

McCrone, J. T., Woods, R. J., Monto, A. S., Martin, E. T., and Lauring, A. S. 2020. The effective population size and mutation rate of influenza a virus in acutely infected individuals.

Murcia, P. R., Baillie, G. J., Daly, J., Elton, D., Jervis, C., Mumford, J. A., Newton, R., Parrish, C. R., Hoelzer, K., Dougan, G., Parkhill, J., Lennard, N., Ormond, D., Moule, S., Whitwham, A., McCauley, J. W., McKinley, T. J., Holmes, E. C., Grenfell, T., and Wood, J. L. N. 2010. Intra- and interhost evolutionary dynamics of equine influenza virus. J. Virol., 84(14): 6943–6954.

Partlow, E. A., Jaeggi-Wong, A., Planitzer, S. D., Berg, N., Li, Z., and Ivanovic, T. 2025. Influenza a virus rapidly adapts particle shape to environmental pressures. Nat. Microbiol., 10(3): 784–794.

Pawelek, K. A., Huynh, G. T., Quinlivan, M., Cullinane, A., Rong, L., and Perelson, A. S. 2012. Modeling within-host dynamics of influenza virus infection including immune responses. PLoS Comput. Biol., 8(6): e1002588.

Pervolaraki, K., Rastgou Talemi, S., Albrecht, D., Bormann, F., Bamford, C., Mendoza, J. L., Garcia, K. C., McLauchlan, J., Höfer, T., Stanifer, M. L., and Boulant, S. 2018. Differential induction of interferon stimulated genes between type I and type III interferons is independent of interferon receptor abundance. PLoS Pathog., 14(11): e1007420.

Phipps, K. L., Ganti, K., Jacobs, N. T., Lee, C.-Y., Carnaccini, S., White, M. C., Manandhar, M., Pickett, B. E., Tan, G. S., Ferreri, L. M., Perez, D. R., and Lowen, A. C. 2020. Collective interactions augment influenza a virus replication in a host-dependent manner. Nat. Microbiol., 5(9): 1158–1169.

Plante, J. A., Liu, Y., Liu, J., Xia, H., Johnson, B. A., Lokugamage, K. G., Zhang, X., Muruato, A. E., Zou, J., Fontes-Garfias, C. R., Mirchandani, D., Scharton, D., Bilello, J. P., Ku, Z., An, Z., Kalveram, B., Freiberg, A. N., Menachery, V. D., Xie, X., Plante, K. S., Weaver, S. C., and Shi, P.-Y. 2021. Spike mutation D614G alters SARS-CoV-2 fitness. Nature, 592(7852): 116–121.

Rabouw, H. H., Schokolowski, J., Müller, M., Baars, M. J. D., Dost, A. F. M., Bestebroer, T. M., Püschel, J., Clevers, H., Fouchier, R. A. M., and Tanenbaum, M. E. 2026. Live-cell single-vRNP imaging identifies viral gene expression signatures that shape influenza infection heterogeneity. Cell Syst., 17(2): 101489.

Rayner, C. R., Chanu, P., Gieschke, R., Boak, L. M., and Jonsson, E. N. 2008. Population pharmacokinetics of oseltamivir when coadministered with probenecid. J. Clin. Pharmacol., 48(8): 935–947.

Reffsin, S., Miller, J., Ayyanathan, K., Dunagin, M. C., Heyman, Y., Jain, N., Schultz, D. C., Cherry, S., and Raj, A. 2026. Single-cell susceptibility to viral infection is driven by variable cell states. Cell, 189(1): 179–195.e21.

Rice, P., Martin, E., He, J.-R., Frank, M., DeTolla, L., Hester, L., O’Neill, T., Manka, C., Benjamin, I., Nagarsekar, A., Singh, I., and Hasday, J. D. 2005. Febrile-range hyperthermia augments neutrophil accumulation and enhances lung injury in experimental gram-negative bacterial pneumonia. J. Immunol., 174(6): 3676–3685.

Rivera-Cardona, J., Mahajan, T., Thayer, E. A., Kakuturu, N. R., Teo, Q. W., Lederer, J., Rowland, E. F., Heimburger, K., Sun, J., McDonald, C. A., Mickelson, C. K., Langlois, R. A., Wu, N. C., Milenkovic, O., Maslov, S., and Brooke, C. B. 2026. Intrinsic OASL expression governs heterogeneity in interferon induction during influenza a virus infection. Proc. Natl. Acad. Sci. U. S. A., 123(1): e2509560123.

Roach, S. N., Shepherd, F. K., Mickelson, C. K., Fiege, J. K., Thielen, B. K., Pross, L. M., Sanders, A. E., Mitchell, J. S., Robertson, M., Fife, B. T., and Langlois, R. A. 2024. Tropism for ciliated cells is the dominant driver of influenza viral burst size in the human airway. Proc. Natl. Acad. Sci. U. S. A., 121(31): e2320303121.

Robinot, R., Hubert, M., de Melo, G. D., Lazarini, F., Bruel, T., Smith, N., Levallois, S., Larrous, F., Fernandes, J., Gellenoncourt, S., Rigaud, S., Gorgette, O., Thouvenot, C., Trébeau, C., Mallet, A., Duménil, G., Gobaa, S., Etournay, R., Lledo, P.-M., Lecuit, M., Bourhy, H., Duffy, D., Michel, V., Schwartz, O., and Chakrabarti, L. A. 2021. SARS-CoV-2 infection induces the dedifferentiation of multiciliated cells and impairs mucociliary clearance. Nat. Commun., 12(1): 4354.

Russell, A. B., Trapnell, C., and Bloom, J. D. 2018. Extreme heterogeneity of influenza virus infection in single cells. Elife, 7(e32303).

Saenz, R. A., Quinlivan, M., Elton, D., Macrae, S., Blunden, A. S., Mumford, J. A., Daly, J. M., Digard, P., Cullinane, A., Grenfell, B. T., McCauley, J. W., Wood, J. L. N., and Gog, J. R. 2010. Dynamics of influenza virus infection and pathology. J. Virol., 84(8): 3974–3983.

Scherer, E. M., Babiker, A., Adelman, M. W., Allman, B., Key, A., Kleinhenz, J. M., Langsjoen, R. M., Nguyen, P.-V., Onyechi, I., Sherman, J. D., Simon, T. W., Soloff, H., Tarabay, J., Varkey, J., Webster, A. S., Weiskopf, D., Weissman, D. B., Xu, Y., Waggoner, J. J., Koelle, K., Rouphael, N., Pouch, S. M., and Piantadosi, A. 2022. SARS-CoV-2 evolution and immune escape in immunocompromised patients. N. Engl. J. Med., 386(25): 2436–2438.

Seki, M., Yanagihara, K., Higashiyama, Y., Fukuda, Y., Kaneko, Y., Ohno, H., Miyazaki, Y., Hirakata, Y., Tomono, K., Kadota, J., Tashiro, T., and Kohno, S. 2004. Immunokinetics in severe pneumonia due to influenza virus and bacteria coinfection in mice. Eur. Respir. J., 24(1): 143–149.

Shi, Y. T., Harris, J. D., Martin, M. A., and Koelle, K. 2024. Transmission bottleneck size estimation from DE novo viral genetic variation. Mol. Biol. Evol., 41(1).

Shi, Y. T., Martin, M. A., Weissman, D. B., and Koelle, K. 2026. Genetic drift acts strongly on influenza virus populations within acute human infections but is obscured by other factors within acutely infected swine. Virus Evol., 12(1).

Shinya, K., Ebina, M., Yamada, S., Ono, M., Kasai, N., and Kawaoka, Y. 2006. Avian flu: influenza virus receptors in the human airway. Nature, 440(7083): 435–436.

Shoemaker, L. G., Sullivan, L. L., Donohue, I., Cabral, J. S., Williams, R. J., Mayfield, M. M., Chase, J. M., Chu, C., Harpole, W. S., Huth, A., HilleRisLambers, J., James, A. R. M., Kraft, N. J. B., May, F., Muthukrishnan, R., Satterlee, S., Taubert, F., Wang, X., Wiegand, T., Yang, Q., and Abbott, K. C. 2020. Integrating the underlying structure of stochasticity into community ecology. Ecology, 101(2): e02922.

Sjaastad, L. E., Fay, E. J., Fiege, J. K., Macchietto, M. G., Stone, I. A., Markman, M. W., Shen, S., and Langlois, R. A. 2018. Distinct antiviral signatures revealed by the magnitude and round of influenza virus replication in vivo. Proc. Natl. Acad. Sci. U. S. A., 115(38): 9610–9615.

Smith, A. M. and Perelson, A. S. 2011. Influenza a virus infection kinetics: quantitative data and models. Wiley Interdiscip. Rev. Syst. Biol. Med., 3(4): 429–445.

Smith, C. M., Kulkarni, H., Radhakrishnan, P., Rutman, A., Bankart, M. J., Williams, G., Hirst, R. A., Easton, A. J., Andrew, P. W., and O’Callaghan, C. 2014. Ciliary dyskinesia is an early feature of respiratory syncytial virus infection. Eur. Respir. J., 43(2): 485–496.

Sobel Leonard, A., McClain, M. T., Smith, G. J. D., Wentworth, D. E., Halpin, R. A., Lin, X., Ransier, A., Stockwell, T. B., Das, S. R., Gilbert, A. S., Lambkin-Williams, R., Ginsburg, G. S., Woods, C. W., and Koelle, K. 2016. Deep sequencing of influenza a virus from a human challenge study reveals a selective bottleneck and only limited intrahost genetic diversification. J. Virol., 90(24): 11247–11258.

Steele, J. H. 1985. A comparison of terrestrial and marine ecological systems. Nature, 313(6001): 355–358.

Tataru, P., Bataillon, T., and Hobolth, A. 2015. Inference under a Wright-Fisher model using an accurate beta approximation. Genetics, 201(3): 1133–1141.

Tonkin-Hill, G., Martincorena, I., Amato, R., Lawson, A. R. J., Gerstung, M., Johnston, I., Jackson, D. K., Park, N., Lensing, S. V., Quail, M. A., Gonçalves, S., Ariani, C., Spencer Chapman, M., Hamilton, W. L., Meredith, L. W., Hall, G., Jahun, A. S., Chaudhry, Y., Hosmillo, M., Pinckert, M. L., Georgana, I., Yakovleva, A., Caller, L. G., Caddy, S. L., Feltwell, T., Khokhar, F. A., Houldcroft, C. J., Curran, M. D., Parmar, S., COVID-19 Genomics UK (COG-UK) Consortium, Alderton, A., Nelson, R., Harrison, E. M., Sillitoe, J., Bentley, S. D., Barrett, J. C., Torok, M. E., Goodfellow, I. G., Langford, C., Kwiatkowski, D., and Wellcome Sanger Institute COVID-19 Surveillance Team 2021. Patterns of within-host genetic diversity in SARS-CoV-2. Elife, 10(e66857).

Ulker, N. and Samuel, C. E. 1987. Mechanism of interferon action. II. induction and decay kinetics of the antiviral state and protein P54 in human amnion U cells treated with gamma interferon. The Journal of biological chemistry, 262(35): 16804–16807.

van Riel, D., den Bakker, M. A., Leijten, L. M. E., Chutinimitkul, S., Munster, V. J., de Wit, E., Rimmelzwaan, G. F., Fouchier, R. A. M., Osterhaus, A. D. M. E., and Kuiken, T. 2010. Seasonal and pandemic human influenza viruses attach better to human upper respiratory tract epithelium than avian influenza viruses. Am. J. Pathol., 176(4): 1614–1618.

VanInsberghe, D., McBride, D. S., DaSilva, J., Stark, T. J., Lau, M. S. Y., Shepard, S. S., Barnes, J. R., Bowman, A. S., Lowen, A. C., and Koelle, K. 2024. Genetic drift and purifying selection shape within-host influenza a virus populations during natural swine infections. PLoS Pathog., 20(4): e1012131.

Varble, A., Albrecht, R. A., Backes, S., Crumiller, M., Bouvier, N. M., Sachs, D., García-Sastre, A., and tenOever, B. R. 2014. Influenza a virus transmission bottlenecks are defined by infection route and recipient host. Cell Host Microbe, 16(5): 691–700.

Vasseur, D. A. and Yodzis, P. 2004. The color of environmental noise. Ecology, 85(4): 1146–1152.

Vega, V. L., Rodríguez-Silva, M., Frey, T., Gehrmann, M., Diaz, J. C., Steinem, C., Multhoff, G., Arispe, N., and De Maio, A. 2008. Hsp70 translocates into the plasma membrane after stress and is released into the extracellular environment in a membrane-associated form that activates macrophages. J. Immunol., 180(6): 4299–4307.

Von Magnus, P. 1954. Incomplete forms of influenza virus. Adv. Virus Res., 2: 59–79.

Wallace, L. E., Liu, M., van Kuppeveld, F. J. M., de Vries, E., and de Haan, C. A. M. 2021. Respiratory mucus as a virus-host range determinant. Trends Microbiol., 29(11): 983–992.

Wilker, P. R., Dinis, J. M., Starrett, G., Imai, M., Hatta, M., Nelson, C. W., O’Connor, D. H., Hughes, A. L., Neumann, G., Kawaoka, Y., and Friedrich, T. C. 2013. Selection on haemagglutinin imposes a bottleneck during mammalian transmission of reassortant H5N1 influenza viruses. Nat. Commun., 4(1): 2636.

Zhou, B., Thao, T. T. N., Hoffmann, D., Taddeo, A., Ebert, N., Labroussaa, F., Pohlmann, A., King, J., Steiner, S., Kelly, J. N., Portmann, J., Halwe, N. J., Ulrich, L., Trüeb, B. S., Fan, X., Hoffmann, B., Wang, L., Thomann, L., Lin, X., Stalder, H., Pozzi, B., de Brot, S., Jiang, N., Cui, D., Hossain, J., Wilson, M. M., Keller, M. W., Stark, T. J., Barnes, J. R., Dijkman, R., Jores, J., Benarafa, C., Wentworth, D. E., Thiel, V., and Beer, M. 2021. SARS-CoV-2 spike D614G change enhances replication and transmission. Nature, 592(7852): 122–127.

